# SARS-CoV-2 hijacks fragile X mental retardation proteins for efficient infection

**DOI:** 10.1101/2023.09.01.555899

**Authors:** Dimitriya H. Garvanska, Rojelio E. Alvarado, Filip Oskar Mundt, Emma Nilsson, Josephine Kerzel Duel, Fabian Coscia, Richard Lindqvist, Kumari Lokugamage, Bryan A. Johnson, Jessica A. Plante, Dorothea R. Morris, Michelle N. Vu, Leah K. Estes, Alyssa M. McLeland, Jordyn Walker, Patricia A. Crocquet-Valdes, Blanca Lopez Mendez, Kenneth S. Plante, David H. Walker, Melanie Bianca Weisser, Anna K. Overby, Matthias Mann, Vineet D. Menachery, Jakob Nilsson

## Abstract

Viruses interact with numerous host factors to facilitate viral replication and to dampen antiviral defense mechanisms. We currently have a limited mechanistic understanding of how SARS-CoV-2 binds host factors and the functional role of these interactions. Here, we uncover a novel interaction between the viral NSP3 protein and the fragile X mental retardation proteins (FMRPs: FMR1 and FXR1-2). SARS-CoV-2 NSP3 mutant viruses preventing FMRP binding have attenuated replication *in vitro* and have delayed disease onset *in vivo*. We show that a unique peptide motif in NSP3 binds directly to the two central KH domains of FMRPs and that this interaction is disrupted by the I304N mutation found in a patient with fragile X syndrome. NSP3 binding to FMRPs disrupts their interaction with the stress granule component UBAP2L through direct competition with a peptide motif in UBAP2L to prevent FMRP incorporation into stress granules. Collectively, our results provide novel insight into how SARS-CoV-2 hijacks host cell proteins for efficient infection and provides molecular insight to the possible underlying molecular defects in fragile X syndrome.

## Introduction

Viruses encode a limited number of proteins and are thus highly dependent on interactions with cellular host factors to efficiently replicate in host cells [1, 2]. Furthermore, viruses dampen innate immune signaling by interfering with distinct steps in the cellular signaling cascades. A common target of viruses is to disrupt the formation of stress granules which have been implicated as signaling hubs for antiviral signaling [3–6]. Stress granules, large membrane less protein-RNA assemblies, form in the cytoplasm in response to various stress signals including viral infections [7]. Induced by host translation inhibition, stress granules are known to sequester RNA. However, they are also composed of a large number of RNA-binding proteins including G3BP1/2 and UBAP2L that play a key role in nucleating and coordinating stress granule formation [8–11]. Importantly, over 250 host proteins have been implicated in playing a role during stress granule formation, highlighting the complexity of this process [9].

Given the link to antiviral defenses, viruses have developed strategies to disrupt stress granule formation and even hijacked these factors to facilitate their replication [12, 13]. For coronaviruses, viral proteins have been implicated in disrupting stress granule formation including the nucleocapsid protein (N) and accessory ORFs [14–16]. Indeed, we and others recently showed that the N protein of SARS-CoV-2 and SARS-CoV contains an ΦxFG motif that binds to G3BP1/2 to disrupt stress granule formation [17–20]. This represents one of several mechanisms SARS-CoV-2 uses to antagonize cellular antiviral mechanisms [21, 22]. Similarly, members of both the old and new world alphaviruses bind G3BP1/2 proteins through ΦxFG motifs in the hypervariable domains of the NSP3 protein to facilitate viral replication complex assembly and disruption of stress granule formation [23, 24]. Interestingly, eastern equine encephalitis virus (EEEV) binds to both G3BP1/2 and the FMR1/FXR1-2 (FMRPs) through distinct sequences in their hypervariable domain [25, 26]. FMRPs, RNA binding proteins, are also recruited to stress granules and deregulated expression of or mutations in FMR1 results in fragile X syndrome, the most common form of inherited mental retardation [27–30]. The exact underlying molecular cause of fragile X syndrome is still not fully understood, but mutations in UBAP2L and G3BP1/2 have also been linked to mental retardation, highlighting the importance of stress granules to host functions [31].

In this manuscript, we uncover a novel direct interaction between SARS-CoV-2 NSP3 and FMRPs and uncover its role during viral infection in molecular detail.

## Results

### The SARS-CoV-2 NSP3 protein binds to FMRPs

To understand SARS-CoV-2 interactions with cellular host factors, we focused on the multifunctional ∼200 kDa NSP3 protein [32]. NSP3, the largest coronavirus protein, has multiple domains with functions essential for viral infection. While there is significant variation across the coronavirus (CoV) family, eight domains are common to all coronavirus NSP3 proteins including two ubiquitin-like domains, the hypervariable region, macrodomain, papain-like protease, zinc-finger domain, and the two Y domains of unknown function (Fig. 1A). Notably, NSP3 multimers also form pore like structures in double-membrane vesicles housing the CoV replication complex [33]. Given the large size, transmembrane domains, and critical roles during infection, NSP3 has been difficult to study, and its many functions remain poorly understood.

**Figure 1.**
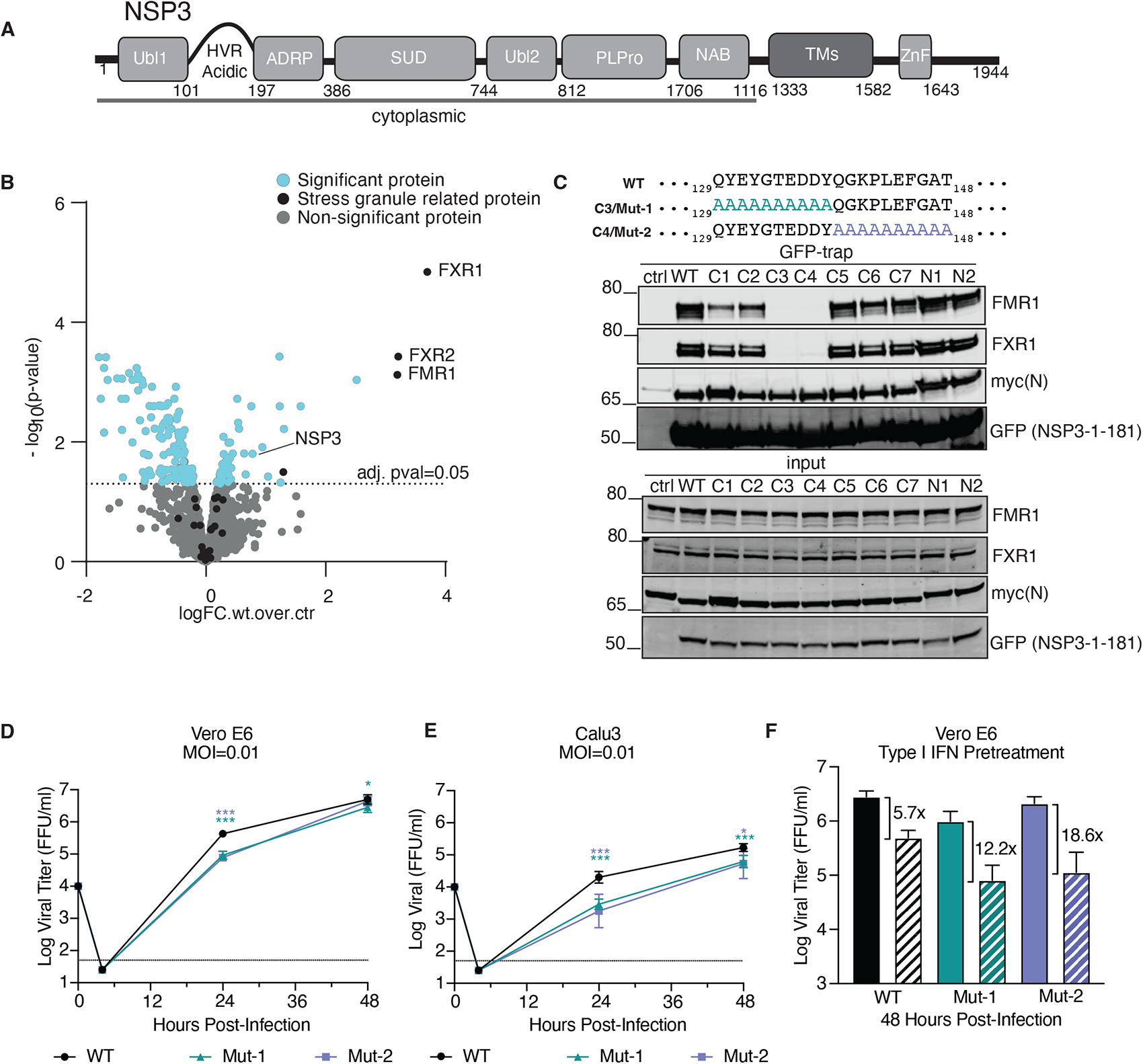
An NSP3 – FMRPs interaction is required for efficient SARS-CoV-2 replication. A) Schematic of NSP3 protein with distinct domains indicated. B) Interactome of the cytoplasmic domains of NSP3 in HeLa cells. Data from 4 technical replicates. C) Interaction of NSP3 mutants with FXR1, FMR1 and myc-tagged N protein to map binding sites. Each variant has 10 amino acids mutated to Alanine. Representative of at 2 independent experiments. D-E) Vero E6 cells or Calu3 cells were infected with the indicated SARS-CoV-2 viruses and viral titers measured at 24 and 48 hours post infection (n=6). F) Vero E6 cells were pretreated with control (solid) or 100 unit of type I IFN (hashed) for 16 hours and then infected with the indicated SARS-CoV-2 viruses and viral titers measured after 48 hours (n=6). Statistical analysis measured by two-tailed Student’s t-test: ***p<0.001, *p<0.05.

For this reason, we chose to explore SARS-CoV-2 NSP3’s interaction with host factors. The pore structure indicates that the N-terminal portion of NSP3 protrudes into the cytoplasm and likely facilitates numerous interactions with host proteins [33]. Therefore, we expressed, and affinity purified a YFP tagged version of the cytosolic part of NSP3 and compared the associated proteins to a control purification using quantitative label free mass spectrometry. Strikingly, the most prominent cellular host factors co-purifying with NSP3 was the RNA binding protein FMR1 and the highly related FXR1 and FXR2 proteins (FMR1, FXR1 and FXR2 collectively referred to here as FMRPs) (Fig. 1B and Supplemental Table 1). The NSP3-FMRP interaction has also been noted in other high throughput screens of SARS-CoV-2 proteins, but its role in viral infection is unknown [34, 35].

To determine which region of NSP3 is interacting with FMRPs, we immunopurified a panel of YFP tagged NSP3 fragments expressed in HeLa cells and monitored binding to endogenous FXR1 by western blotting. Our results revealed that the N-terminal 181 amino acids of NSP3 were capable of binding FXR1 (Supplemental Fig. S1A). Due to its large size, multiple domains, and sequence diversity, it is difficult to compare NSP3 similarity across the coronavirus family. To determine if the observed NSP3/FMRP interaction occurs in other CoVs, we generated N-terminal fragments of five human coronaviruses outside the Sarbeco coronaviruses. However, immunopurifications with these N-terminal regions of NSP3 from these other coronaviruses revealed that tight binding of NSP3 to FMRPs was restricted to SARS-CoV-2 (Supplemental Fig. S1B).

To more precisely identify the binding site of FMRPs we mutated blocks of 10 amino acids to Ala in the intrinsically disordered regions of NSP3 1-181. In total we generated 9 mutants with 2 mutants covering the intrinsically disordered N-terminus and 7 the intrinsically disordered region following the Ubl1 domain. These constructs where expressed in cells together with the viral N protein and immunopurified. Our results established that binding to FMRPs was mediated by a stretch of 20 amino acids in the hypervariable region (HVR) following the Ubl1 domain (Fig. 1C). Notably, mutation of this region did not affect the NSP3-N interaction in agreement with predictions from the structure of the complex [36]. While there is variation in the HVR, this 20 amino acid sequence was conserved in all of the Sarbeco family of coronaviruses (Supplemental Fig. S1C); no similar sequence motifs were found in other coronaviruses. Together, our finding defines the motif in NSP3 binding to FMRPs and indicate its conservation across the Sarbeco virus family.

### SARS-CoV-2 NSP3 mutant viruses have attenuated replication *in vitro*

Having established an interaction between NSP3 and FMRPs within a 20 amino acid region in the HVR, we next sought to determine the impact of this interaction on SARS-CoV-2 infection. Utilizing our reverse genetic system [37, 38], we generated two recombinant SARS-CoV-2 mutant viruses (mut1 and mut2) with the 10 alanine mutations in NSP3 preventing FMRP binding (C3 and C4 in Fig. 1C). Both NSP3 mutants were recovered with normal stock titers and plaque morphology. Examining VeroE6 cells, the NSP3 mutant viruses were attenuated in replication 24 hours post infection (HPI) relative to the WT SARS-CoV-2 (Fig. 1D). While attenuation was mostly absent at 48 HPI, the reduced capacity of the NSP3 mutants in the type I interferon (IFN) deficient VeroE6 was noteworthy. We subsequently examined replication in Calu3 cells, an IFN responsive respiratory cell line (Fig. 1E). Similar to VeroE6 cells, NSP3 mut1 and mut2 were attenuated compared to WT SARS-CoV-2. Together, the data show that disruption of this 20 amino-acid stretch in NSP3 attenuated SARS-CoV-2 replication *in vitro*.

While type I IFN is a major driver of the antiviral response in cells additional antiviral mechanisms exist. The attenuation of the NSP3 mutants in IFN deficient VeroE6 cells suggests interferon stimulated genes (ISGs) may not drive attenuation. To evaluate IFN sensitivity, we pretreated VeroE6 cells with type I IFN and infected with SARS-CoV-2 WT and NSP3 mutants (Fig. 1F). Following ISG activation, WT viruses had a 0.5 to 1.25 log reduction in viral titer compared to untreated. However, NSP3 mutants had a 2 to 3-fold increase in sensitivity compared to WT. Compared to IFN sensitive SARS-CoV-2 NSP16 mutants (>10,000 fold titer reduction) [39], the modest susceptibility of NSP3 mutants suggest antagonism of an antiviral pathway unrelated to the ISG response, governing most of the attenuation.

### NSP3 mutants are attenuated during early *in vivo* infection

To investigate the role of the NSP3-FMRP interaction *in vivo*, we next infected 3-to-4 week old hamsters with WT and NSP3 mutant SARS-CoV-2 evaluating weight loss and disease over 7 days (Supplemental Fig. S2A). In addition, animals were nasal washed, euthanized and tissue collected for further analysis. Hamsters infected with NSP3 mut1 and mut2 showed no significant change in weight loss or disease relative to WT control animals (Fig. 2A). Similarly, we detected no or little significant changes in viral titers in either the nasal wash or lungs infected with NSP3 mutants as compared to WT (Fig. 2B-C). However, despite similar viral titers, histopathology results indicate early attenuation of the NSP3 mutants. Examining antigen staining, WT SARS-CoV-2 infected hamsters show robust viral staining throughout the lung parenchyma and airways on day 2 post infection (Fig. 2D and Supplemental Fig. S2B). In contrast, both NSP3 mutants had significantly less antigen staining at day 2 in the lungs (Fig. 2E-G and Supplemental Fig. 2B). While antigen staining rebounded at day 4 to similar levels as WT, the results suggest that the NSP3 mutants had reduced and delayed infection *in vivo*. Similarly, H&E histopathology shows a significant reduction in cellular infiltration and damage in NSP3 mutant infected hamsters as compared to control (Fig. 2H). WT SARS-CoV-2 infected lungs showed wide-spread damage with multifocal interstitial pneumonia, perivasculitis, bronchiolitis, and peribronchiolitis 2 days post infection. In contrast, both NSP3 mutants had more focal disease with less extensive damage at day 2. While later timepoints showed similar histolopathological lesions, the results indicate the loss of NSP3/FMRP binding delays SARS-CoV-2 disease induction.

**Figure 2.**
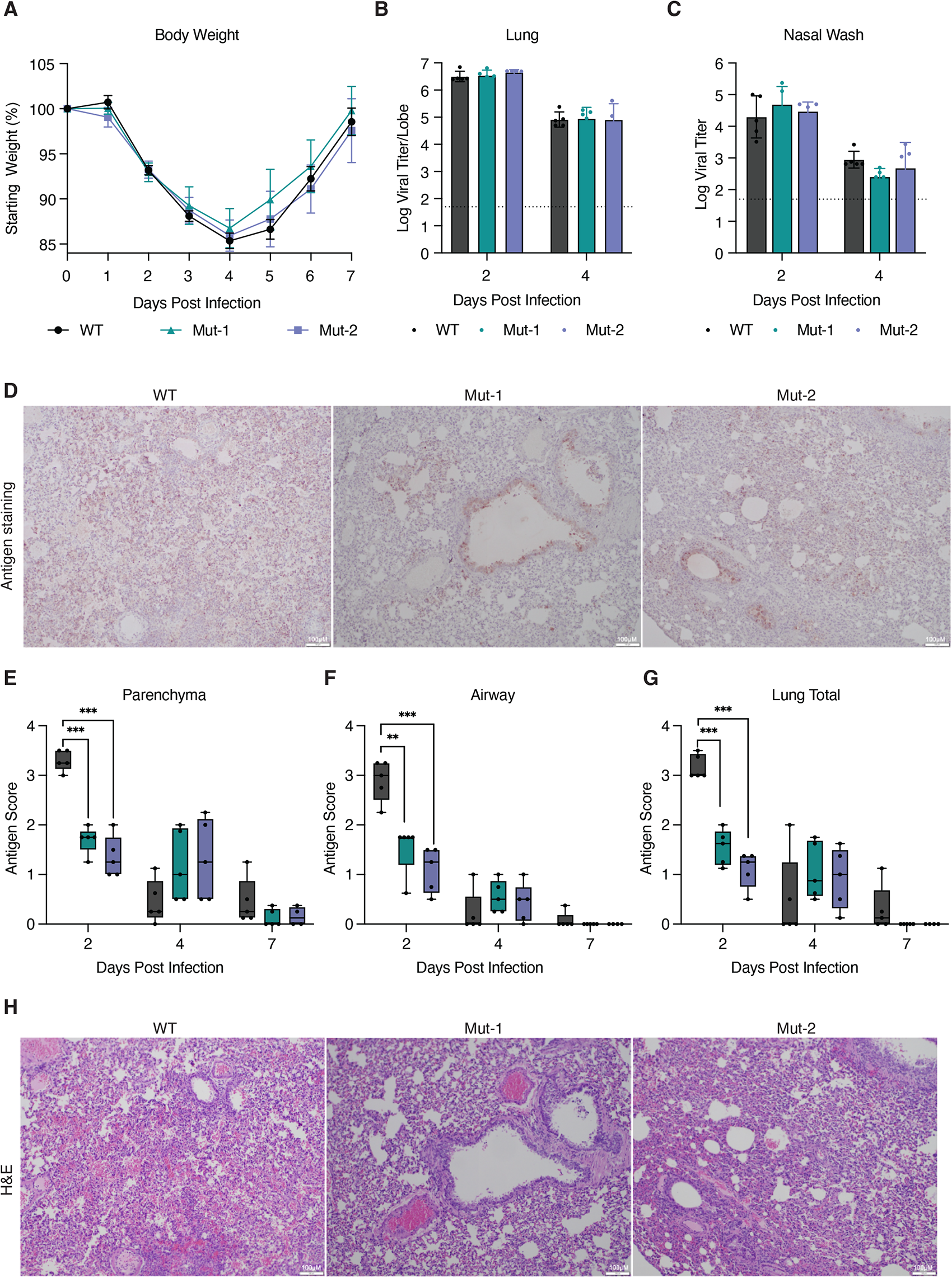
*In vivo* characterization of SARS-CoV-2 virus unable to bind FMRPs. A) Golden Syrian hamsters were infected with 10^5^ plaque forming units (PFU) of WT SARS-CoV-2 (n=15), NSP3 mutants (n=15), or mock (PBS, n=15) and monitored for weight loss and signs of disease over a 7 day time course. B-C) At days 2 and 4 post infection, infected hamsters (n=5) were nasal washed and subsequently euthanized and tissue collected to assay viral titers from B) lung or C) nasal wash. D) Lung tissue sections were stained for viral antigen (nucleocapsid) at day 2 for WT and NSP3 mutants infected animals. E-G) Antigen staining was scored in the parenchyma, airway, and by total in a blinded manner. Data are mean showing minimum and maximum (n=5). Statistical analysis measured by two-tailed Student’s t-test: ***p<0.001, **p<0.01. H) Parallel lung tissue sections demonstrated more severe lesions at day 2 post infection for WT infected hamsters versus NSP mutant infected animals.

### NSP3 binds the RNA binding region of the FMRP KH domains

To understand how the NSP3-FMRP complex contributes to viral infection we first aimed at obtaining a detailed molecular and structural understanding of the complex. FMRPs are composed of distinct domains, and we therefore constructed a panel of tagged FXR1 fragments and monitored binding to NSP3 1-181 by immunoprecipitation (Fig. 3A). Our results revealed that FXR1 1-215 did not bind NSP3 while FXR1 1-360 did, suggesting that the two central KH domains mediate binding. Indeed FXR1 215-360 comprising the two central KH domains was sufficient for binding to NSP3 1-181. The central KH domains are almost identical in sequence among the FMRPs explaining why we observe all FMRPs binding to NSP3. Interestingly, the fragile X syndrome disease mutation, I304N, fully blocked the interaction between FXR1 and NSP3 (Supplemental Fig. S3A).

**Figure 3.**
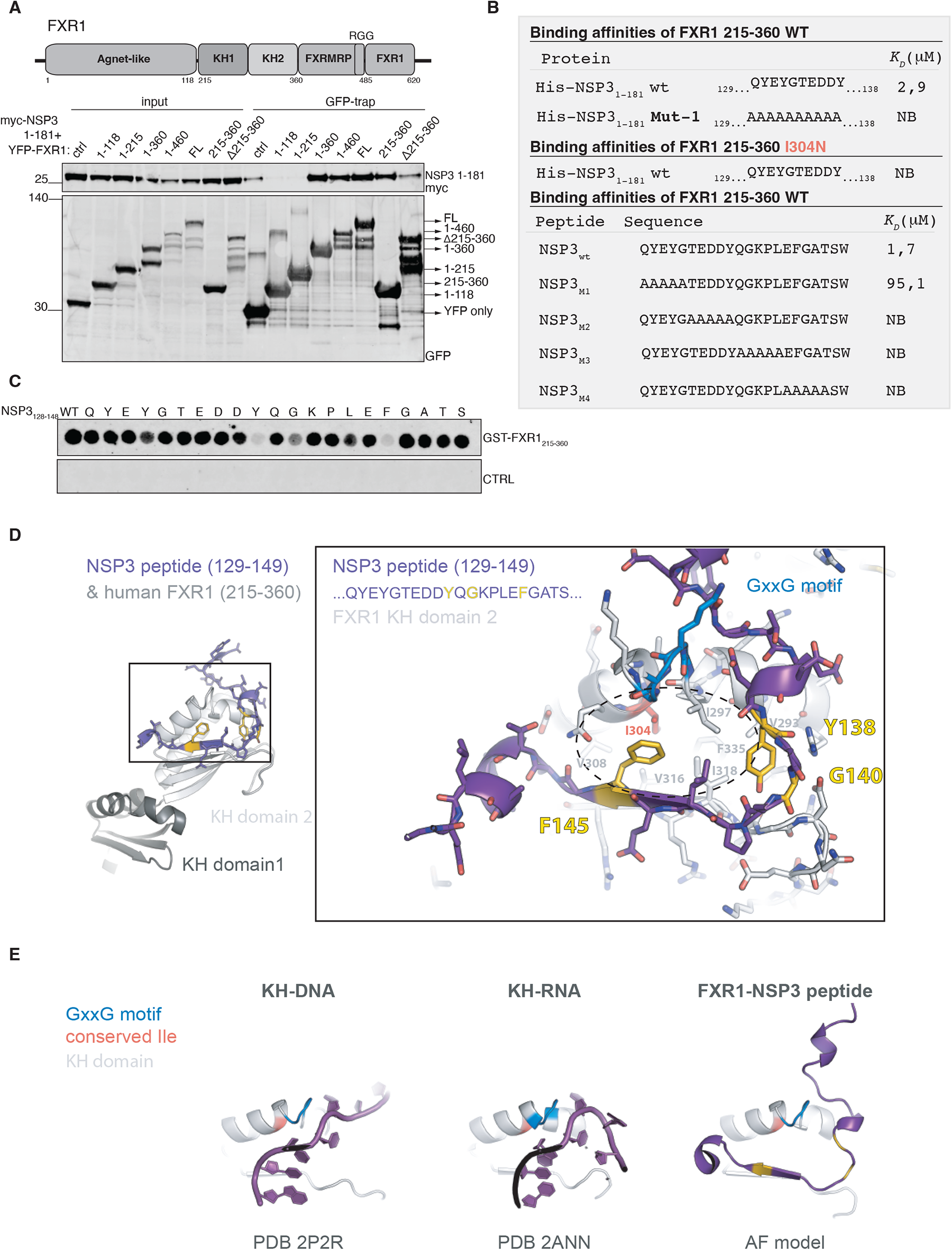
NSP3 binds to the FMRP KH domains similar to how RNA binds. A) Schematic of FXR1 and immunopurification of the indicated FXR1 fragments and binding to NSP3 1-181 determined. Representative of 2 independent experiments. B) Affinity measurements by ITC of the indicated proteins and peptides. C) Spot array of the indicated NSP3 peptide incubated with purified FXR1 215-360 to map critical residues required for binding. Representative of 2 independent experiments. D) AlphaFold model of the FXR1-NSP3 complex with critical residues in NSP3 indicated by yellow and the I304 residue in FXR1 highlighted. E) Comparison between KH-DNA and KH-RNA structures and the model of FXR1-NSP3.

To investigate if the interaction is direct, we expressed and purified a number of FXR1 fragments as well as NSP3 1-181 WT and mut1 mutant. In size-exclusion chromatography experiments we observed complex formation of FXR1 215-360 and NSP3 1-181 WT (Supplemental Fig. S3B). We measured the affinity between a number of recombinant FXR1 fragments and recombinant NSP3 1-181 using isothermal titration calorimetry (ITC) (Supplemental Fig. S3C). Consistent with our cellular co-purification experiments, we detected specific binding between NSP3 1-181 and FXR1 215-360 measuring the affinity to 2,3-2,9 uM (Fig. 3B and Supplemental Fig. S3C and Supplemental Table 2). This interaction was abolished by the FXR1 I304N mutation and the NSP3 mut1 mutations consistent with the cellular data (Fig. 3B and Supplemental Table 2). Importantly the 20 amino acid region of NSP3 required for interaction was sufficient for binding to FXR1 215-360 with a similar affinity as NSP3 1-181 (Fig. 3B). Collectively this shows that the 20 amino acid sequence of SARS-CoV-2 NSP3 is required and sufficient for interaction with FMRPs. Notably, we confirmed that a reported 23mer NSP3 peptide from alphaviruses binds FXR1 1-122 arguing that these viruses hijack FMPRs by a distinct mechanism (Supplemental Fig. S3C)[25].

We next sought to define the key residues in NSP3 mediating binding to FMRPs. A five amino acid alanine scan through the SARS-CoV-2 NSP3 peptide did not further narrow down the interaction which argues that multiple residues in this region bind FXR1 (Fig. 3B). We therefore conducted a peptide array experiment where we changed single amino acids to alanine residues in the NSP3 20mer peptide and monitored binding to FXR1 215-360 (Fig. 3C). This study pinpointed 3 residues in NSP3 critical for binding to FMRPs: Y138, G140, and F145A. We subsequently generated a structural model of the complex using AlphaFold multimer (Fig. 3D and Supplemental Fig. S3D for pLDDT value)[40]. Interestingly, this model revealed that the NSP3 peptide interacts with the GxxG motif of the FMRP KH2 domain similarly to how RNA and DNA has been shown to bind KH domains [41] (Fig. 3E). This model further revealed an interaction between FXR1 I304, a residue stabilizing the hydrophobic core of the KH domain, and NSP3 F145 providing an explanation for why mutation of these residues abolish binding. Similarly, Y138 and G140 may play a role in stabilizing a NSP3 loop facilitating further interactions with FMRPs. We have been unable to detect direct binding of RNA to the FXR1 KH domains using a reported RNA that binds full length FMR1 [42] (Supplemental Fig. S4A and Supplemental Table 2) suggesting that these KH domains might be preferentially involved in protein-protein interactions rather than RNA binding. Collectively, our results reveal the key residues in SARS-CoV-2 NSP3 protein that bind to the two central KH domains of FMRPs and suggest RNA mimicry by this peptide.

### NSP3 disrupts the interaction between FMRPs and UBAP2L

To get insight into how the NSP3-FMRP interaction antagonizes antiviral host mechanisms, we determined if NSP3 binding rewires the interactome of FMRPs. Using lysate from cells expressing YFP-FXR1, we added either WT NSP3 peptide or a non-binding control peptide (NSP3_m2_ in Fig. 3B) and then affinity purified FXR1. We then use mass spectrometry-based proteomics to quantitatively compare the samples. Our results revealed a striking displacement of stress granule components from FXR1 in the presence of the NSP3 WT peptide (Fig. 4A and Supplemental Table 1). UBAP2L, one of the most affected proteins, shows a strong reduction in co-purification consistent with a possible direct interaction between UBAP2L and FXR1 215-360 shown by a prior two-hybrid screen [43]. To confirm, we conducted the inverse experiment and immunopurified UBAP2L-Venus in the presence of WT NSP3 or mutant NSP3 peptide revealing that the entire FMRP-TDRD3-TOP3B complex was displaced (Fig. 4B and Supplemental Table 1). Consistent with the FXR1 I304N mutant being unable to bind the NSP3 peptide this mutant was also defective in binding to UBAP2L and stress granule components in cells (Supplemental Fig. S4B-C and Supplemental Table 1). A recently reported FXR1 mutant unable to form cytoplasmic granules, FXR1 L351P, also failed to bind UBAP2L [44](Supplemental Fig S4C). Together, the results argue that NSP3 binding disrupts the interactions between UBAP2L and FMRPs.

**Figure 4.**
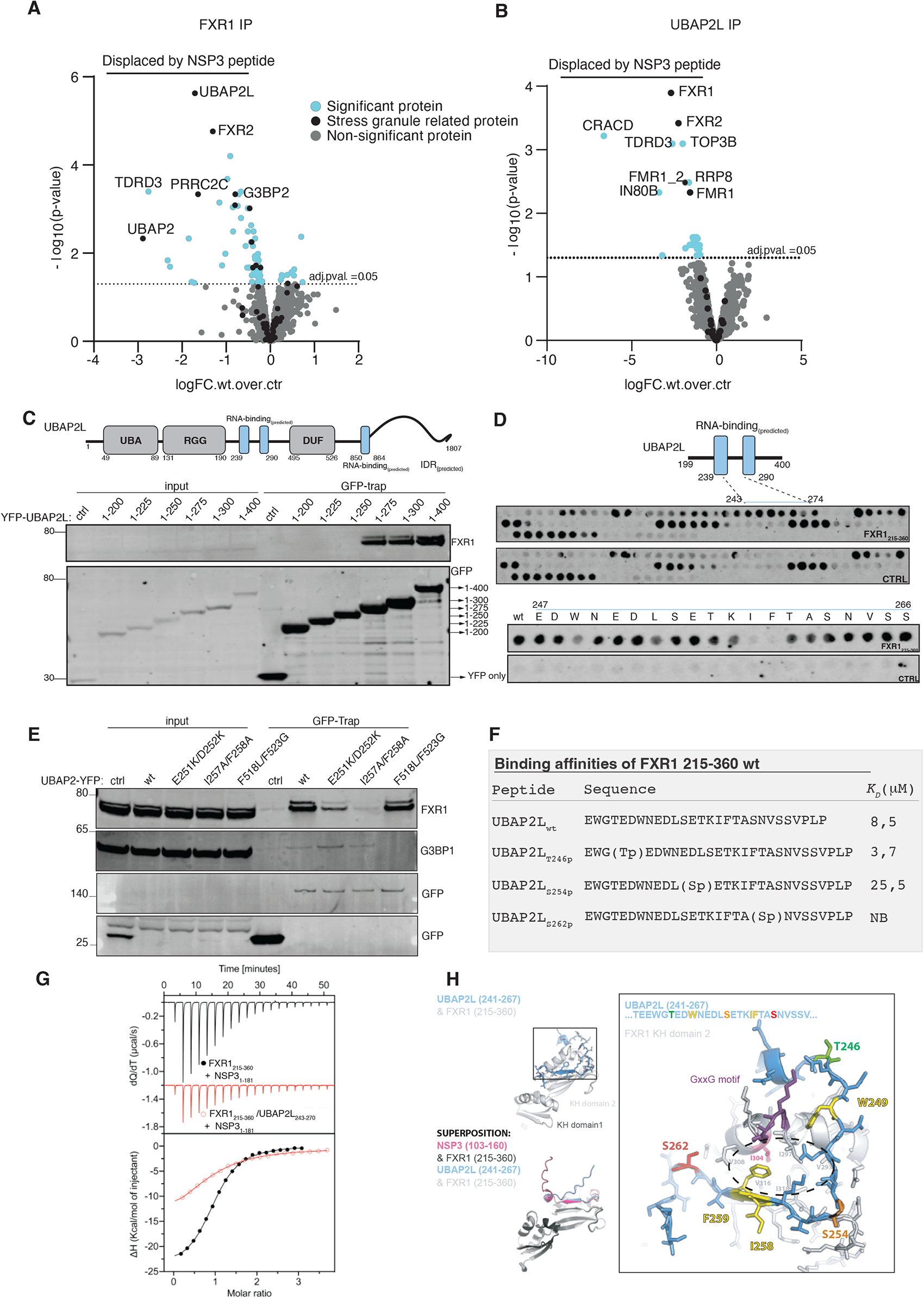
NSP3 disrupts the UBAP2L-FMRP complex. A) FXR1 was affinity purified and incubated with either WT NSP3 or mutant NSP3 peptide and interactomes determined by MS to determined proteins specifically displaced by WT NSP3. Data from 4 technical repeats. B) As A) but using UBAP2L as a bait. Data from 4 technical repeats. C) Schematic of UBAP2L and truncation analysis to identify FXR1 binding site (n=1). D) Peptide array of UBAP2L 199-400 to identify FXR1 binding region and lower part single Ala scan through UBAP2L 247-266 to identify critical residues (n=1). E) Immunopurification of indicated UBAP2L constructs to determine binding to FXR1 and G3BP1. Representative of 2 experiments. F) ITC measurements of indicated UBAP2L peptides to FXR1 215-360. G) Competition between NSP3 peptide and UBAP2L peptide for binding to FXR1 215-360. The black trace is NSP3 binding to FXR1 while the red trace is NSP3 binding to FXR1 preincubated with UBAP2L peptide. H) Alphafold model of the FXR1-UBAP2L complex highlighting critical residues in yellow and phosphorylation sites.

The interactome data suggested that UBAP2L and NSP3 compete for binding to a similar interface on FMRPs. To test this directly, we first mapped the binding site in UBAP2L to FMRPs. A truncation analysis of UBAP2L identified the region from 200-400 as required for interaction (Supplemental Fig. S4D). To further map the site of interaction, we generated a peptide array that covered this region of UBAP2L with 20mer peptides shifted by 2 amino acids at a time. We observed specific binding of FXR1 215-360 to peptides spanning 243-274 in UBAP2L; these results were further supported by immunopurification of UBAP2L fragments (Fig. 4C-D and Supplemental Fig. S4D). An alanine scan through UBAP2L 247-266 pinpointed W249, L253, K257, I258 and F259 as critical residues for binding (Fig. 4D). Based on this finding we generated UBAP2L I258A/F259A and in addition a charge swap mutant, UBAP2L E251K/D252K, which both showed a clear reduction in binding to FXR1 in immunopurifications (Fig. 4E). In contrast mutating the G3BP1/2 binding site in UBAP2L (F518L/F523G) did not affect FXR1 binding (Fig. 4E). This region is conserved in UBAP2 arguing that FMRP interaction mode is conserved between UBAP2L and UBAP2. We measured the affinity of a UBAP2L peptide spanning residues 243-270 which revealed an affinity of 8,5 uM and we confirmed that the NSP3 peptide and UBAP2L peptide competed for binding to FXR1 260-315 by ITC (Fig. 4G).

Consistent with this result, an AlphaFold model of the UBAP2L-FXR1 complex revealed a similar mode of interaction of UBAP2L with the KH2 domains as that observed with NSP3 (Fig. 4H and Supplemental Fig. S5A for pLDDT value). In this model, FXR1 I304 interacts with UBAP2L F259 similar to its interaction with NSP3 F145. We noted several reported phosphorylation sites in the region of UBAP2L binding to FMRPs providing a means to regulate the interaction. To investigate this possibility, we measured the binding affinity of three UBAP2L phosphopeptides to FXR1 by ITC. Interestingly, Thr246 phosphorylation resulted in increased affinity with a Kd of 3,5 while phosphorylation of Ser254 and Ser262 disrupted the interaction (Fig. 4F). The effects of these phosphorylations were consistent with the AlphaFold model of the complex (Fig. 4H and Supplemental Fig. S5B). Collectively these data reveal that SARS-CoV-2 NSP3 competes directly with UBAP2L for binding to FMRPs and displaces the FMRP-TDRD3-TOP3B complex from UBAP2L.

### The incorporation of FMRPs into stress granules is disrupted by NSP3

Our data suggested that the ability of NSP3 to antagonize host cell antiviral mechanisms could be through an effect on stress granule composition and assembly. To investigate this, we investigated FXR1 localization during infection in VeroE6 cells. We observed that FXR1 associated with stress granules in cells expressing low levels of the SARS-CoV-2 N protein. As the levels of N increased reflecting later stages of infection, FXR1 localization shifted and was evenly distributed throughout the cytoplasm (Supplemental Fig. S6A-B). We noted that the total levels of FXR1 increased during infection consistent with a previous study [45] (Supplemental Fig. S6C). Since UBAP2L plays an important role in nucleating stress granules, we speculated that NSP3 affected the ability of FXR1 to associate with these structures through competition during infection. Consistent with this idea FXR1 I304N and FXR1 L351P were unable to form stress granules induced by arsenite (Supplemental Fig. S6D). To test this directly, we expressed NSP3 1-181 WT or NSP3 Mut-2 and monitored the ability of endogenous FXR1 to associate with stress granules following their induction by arsenite (Fig. 5A). In line with our interaction data, we observed that FXR1 was impaired in associating with stress granules in the presence of NSP3 WT, but not NSP3 Mut-2. Importantly this effect appeared to be specific to FXR1 as G3BP1 incorporation was not strongly affected by NSP3. Using live cell imaging to monitor the incorporation of YFP-FXR1 and YFP-G3BP1 into stress granules, we observed a strong reduction of FXR1 incorporation when co-expressed with Cherry tagged NSP3 1-181 WT (Supplemental Fig. S7A) contrasting a small reduction with G3BP1. Similarly, we complemented HeLa UBAP2L KO cells [10] with our UBAP2L mutants and analyzed incorporation of FXR1 into stress granules upon arsenite addition. Preventing the interaction between UBAP2L and FMRPs did not affect the ability of UBAP2L to form stress granules in contrast to the UBAP2L mutant unable to bind G3BP1/2 (Supplemental Fig. S7B). Incorporation of FXR1 into stress granules was strongly impaired in the UBAP2L KO cell lines as expected but could be restored by expressing UBAP2L-YFP. However, mutations in UBAP2L preventing FXR1 binding also prevented efficient incorporation of FXR1 into stress granules (Fig. 5C-D). Collectively, our results shows that NSP3 blocks the UBAP2L-FMRP interaction necessary for association of FMRPs with stress granules and this acts to antagonize antiviral defense mechanisms efficiently during early stages of infection.

**Figure 5.**
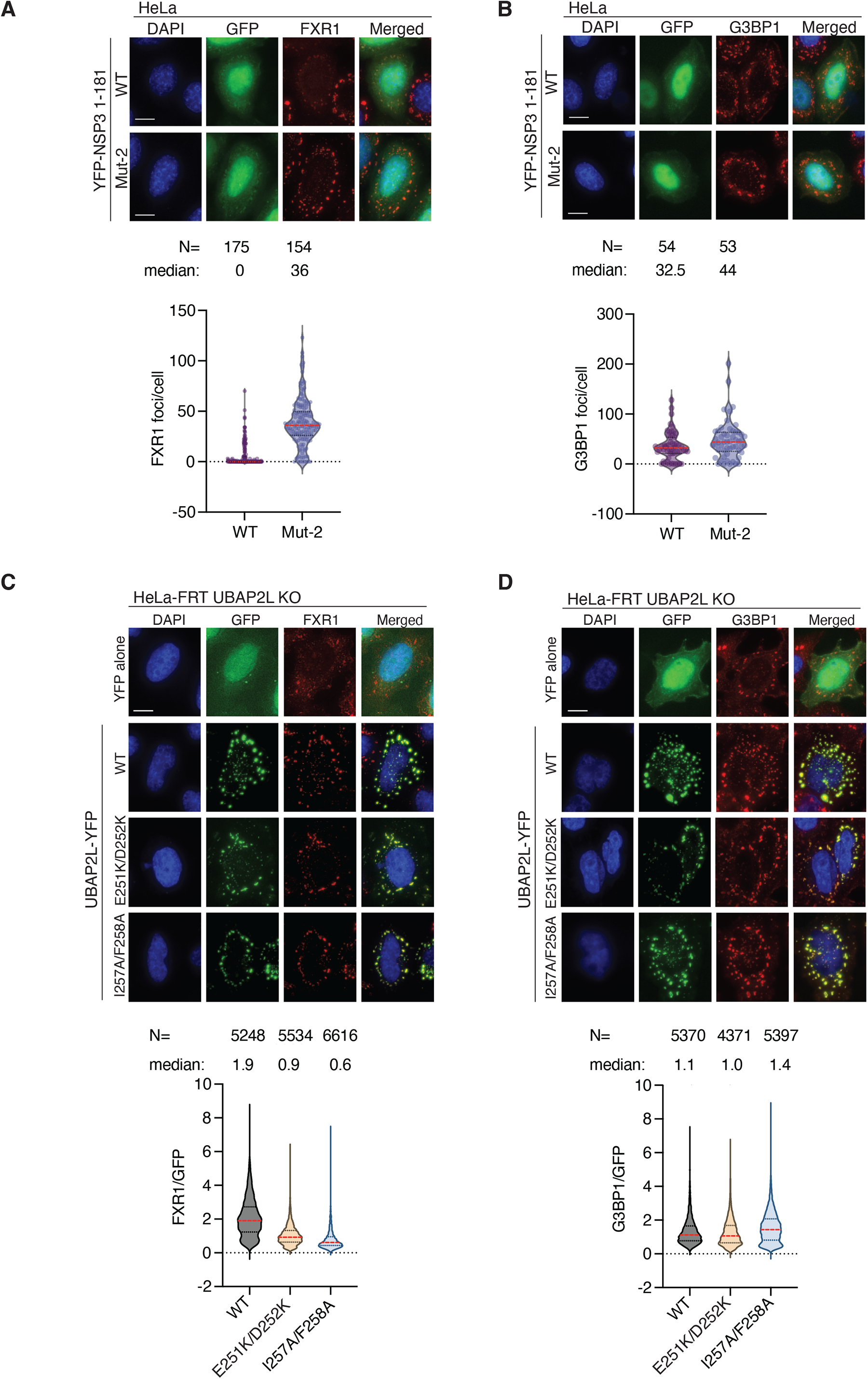
NSP3 prevents incorporation of FMRPs into stress granules. A) Analysis of FXR1 stress granule association in the presence of NSP3 WT or NSP3 A4. HeLa cells were treated 30 minutes with arsenite before fixation and number of FXR1 foci quantified per cell. Only cells expressing NSP3 was analysed. B) Similar as A) but number of G3BP1 foci analysed. C) HeLa UBAP2L KO cells were complemented with UBAP2L-YFP constructs and cells treated with arsenite for 30 minutes before fixation. The fluorescence intensity of FXR1 to UBAP2L (GFP signal) was quantified. D) As in C) but staining for G3BP1. In A-D representative stills from the immunofluorescence is shown with a scale bar of 10 μM indicated in lower left corner. In A-B combined data from 3 experiments is shown in the graphs. In C-D A pool of 4 experiments is shown in the graphs. The median is indicated with red line. The median and number of cells (N) analyzed per condition is indicated above the plot.

## Discussion

Viruses use several strategies to antagonize different aspects of the host antiviral responses. Here, we uncover a novel mechanism by which SARS-CoV-2 NSP3 disrupts the UBAP2L-FMRP interaction to antagonize a specific aspect of stress granule assembly. Specifically, a 20 amino-acid region of the NSP3 hypervariable region, well conserved in Sarbeco coronaviruses, mediates binding to FMRP hereby competing with binding to UBAP2L and thus FMRP incorporation into stress granules. While NSP3 mutants have attenuated virus replication *in vitro* and delayed disease *in vivo*, the impacts are relatively modest. However, NSP3 may blunt parts of stress granule formation early until sufficient N protein ablates the host process later during infection. This function of NSP3 may be even more impactful in susceptible hosts like the aged or immune compromised.

Stress granule formation is a highly complex process with numerous components, cell type specificity, and critical functions in host responses. Importantly, viruses have numerous approaches to disrupt and hijack stress granule elements to facilitate viral infection. The importance of stress granules to the host antiviral response is cemented by the genetic capital SARS-CoV-2 devotes to antagonizing it. For SARS-CoV-2, the N protein uses its ΦxFG motif to prevent G3BP1/2 interactions, powerfully disrupting stress granule formation [18]. In this study, we find NSP3 to be an additional stress granule antagonist in the Sarbecovirus family, disrupting integration of FMRPs via UBAP2L. To our knowledge, SARS-CoV-2 NSP3 is the first example of a virus targeting UBAP2L-FMRP interactions. Interestingly, the required peptide sequence in NSP3 is more extensive than just the ΦxFG found in SARS-CoV-2 N. In our AlphaFold models, both the NSP3 peptide and the UBAP2L peptide engage the FMRP KH domain at a similar site as RNA in reported KH-RNA structures. This is the first example of a KH domain binding to a peptide, suggesting that this protein fold is more versatile in function [41]. If the KH domains of FMRP are binding specific RNAs and whether these are affected by NSP3 are unclear but important to determine. Given that NSP3 binds to N through its Ubl domain, both stress granule disruptive viral proteins would be in close proximity. This viral complex could ensure efficient stress granule disassembly at the sites where nascent viral RNA emerges from the double membrane vesicles through pores formed by NSP3 [33]. Together with the N protein, NSP3 antagonism demonstrates multiple approaches SARS-CoV-2 takes to modulate stress granule functions. Importantly, disruption of either NSP3 and N function on stress granules has implications for SARS-CoV-2 infection and pathogenesis. This may open therapeutic approaches to target these interfaces to facilitate direct antiviral drug treatments and live-attenuated vaccine approaches. However, similar interactions with host proteins increase the possibility of toxicity and off-target effect.

In addition to providing new insight into SARS-CoV-2 biology our work also hints at potential underlying molecular defects of fragile X syndrome. Recent data suggest that brain development is linked to stress granule function, but understanding this will require deeper mechanistic insight into stress granule biology. The molecular detailed understanding of the FMRP-UBAP2L complex described here can potentially help explain its role in brain development and how this is affected by disease mutations such as FMR1 I304N. It will furthermore be interesting to explore if antiviral defense mechanisms are affected in patients with fragile X syndrome or harboring mutations in stress granule components and whether this affects disease trajectory of COVID19.

## Acknowledgements

Work at the Novo Nordisk Foundation Center for Protein Research is supported by grant NNF14CC0001 and JN and MM is supported by a grant from Sygeforsikring Danmark and JN by a grant Independent Research Fund Denmark. We thank the protein production unit and NNF CPR for help with purifying proteins and the medical faculty Umeå University strategic research resource and the Laboratory for Molecular Infection Medicine Sweden for generous support (A.K.Ö.), and the Biochemical Imaging Center at Umeå University and the National Microscopy Infrastructure, NMI (VR-RFI 2016-00968) for assistance in microscopy. Work in AKO lab is supported by the Swedish Research Council (2018-05851). Research at UTMB was supported by grants from NIAID of the United States NIH to (R01-AI153602, R21-AI145400, U19-AI171413, R24-AI120942 to VDM. Research was also supported by STARs Award provided by the University of Texas System to VDM. Trainee funding provided by NIAID of the NIH to MNV (T32-AI060549). We thank Anne-Claude Gingras for providing the HeLa UBAP2L KO cell line.

## Author contributions

DHG conducted all immunopurification, immunofluorescence, live cell and size-exclusion experiments. Viral engineering work that was done by REA, KGL, BAJ and VDM. Viral in vitro and in vivo assays was done by REA, BAJ, JAP, DM, MNV, LKE, AMM, PAC, KSP, DHW. DHW and PAC analyzes pulmonary histopathology. Mass spectrometry analysis was done by FOM, JKD, FC and MM. EN, RL and AO analysed FXR1 incorporation into stress granules during infection. ITC measurements were done by BLM and AlphaFold models by MBW. JN and VDM drafted manuscript.

## Materials and Methods

### Cell Culture for Viral Infections

HeLa, HeLa-FRT parental, HeLa-FRT UBAP2L KO and HEK293 cell lines were cultured in DMEM GlutaMax media (Thermo Fisher Scientific) at 37C in a humidified incubator with 5%CO2. The media was supplemented with 10% FBS (HyClone) and 1% PenSrep (Thermo Fisher Scientific).

Vero E6 cells were grown in high glucose DMEM (Gibco #11965092) with 10% fetal bovine serum and 1x antibiotic-antimycotic. TMPRSS2-expressing Vero E6 cells were grown in low glucose DMEM (Gibco #11885084) with sodium pyruvate, 10% FBS, and 1 mg/mL Geneticin^TM^ (Invitrogen #10131027). Calu-3 2B4 cells were grown in high glucose DMEM (Gibco #11965092) with 10% defined fetal bovine serum, 1 mM sodium pyruvate, and 1x antibiotic-antimycotic.

### Viruses

The SARS-CoV-2 infectious clones were based on the USA-WA1/2020 sequence provided by the World Reference Center of Emerging Viruses and Arboviruses and the USA Centers for Disease Control and Prevention [46]. Mutant viruses (NSP3-Mut1 and NSP3 Mut2) were generated with restriction enzyme-based cloning using gBlocks encoding the mutations (Integrated DNA Technologies) and our reverse genetics system as previously described [37, 47]. Virus stock was generated in TMPRSS2-exrpressing Vero E6 cells to prevent mutations from occurring at the FCS as previously described[48] . Viral RNA was extracted from virus stock and cDNA was generated to verify mutations by Sanger sequencing.

### *In vitro* Infection

Vira infections in Vero E6 and Calu-3 2B4 were carried out as previously described [49]. Briefly, growth media was removed, and cells were infected with WT or mutant SARS-CoV-2 at an MOI of 0.01 for 45 min at 37°C with 5% CO_2_. After absorption, cells were washed three times with PBS and fresh complete growth media was added. Three or more biological replicates were collected at each time point and each experiment was performed at least twice. Samples were titrated with plaque assay or focus forming assays.

### Virus Quantitation via Focus Forming Assay

Focus forming assays (FFAs) were performed as previously described [50]. Briefly, Vero E6 cells were seeded in 96-well plates to be 100% confluent. Samples were serially diluted in serum-free media and 20 µl was used to infect cells. Cells were incubated for 45 min at 37°C with 5% CO_2_ before 0.85% methylcellulose overlay was added. Cells were incubated for 24 h at 37°C with 5% CO_2_. After incubation, overlay was removed, and cells were washed three times with PBS before fixed and virus inactivated by 10% formalin for 30 min at room temperature. Cells were then permeabilized and blocked with 0.1% saponin/0.1% BSA in PBS before incubated with α-SARS-CoV-2 Nucleocapsid primary antibody (Cell Signaling Technology) at 1:1000 in permeabilization/blocking buffer overnight at 4°C. Cells were then washed (3x) with PBS before incubated with Alexa Fluor^TM^ 555-conjugated α-mouse secondary antibody (Invitrogen #A28180) at 1:2000 in permeabilization/blocking buffer for 1 h at room temperature. Cells were washed (3X) with PBS. Fluorescent foci images were captured using a Cytation 7 cell imaging multi-mode reader (BioTek), and foci were counted manually.

### Hamster Infection

Three-to four-week-old male golden Syrian hamsters (HsdHan:AURA strain) were purchased from Envigo. All studies were conducted under a protocol approved by the UTMB Institutional Animal Care and Use Committee and complied with USDA guidelines in a laboratory accredited by the Association for Assessment and Accreditation of Laboratory Animal Care. Procedures involving infectious SARS-CoV-2 were performed in the Galveston National Laboratory ABSL3 facility. Hamsters were intranasally infected with 10^5^ pfu of WT or NSP3 Mut1 or NSP3 Mut2 SARS-CoV-2 in 100 µl. Infected hamsters were weighed and monitored for illness over 7 days. Hamsters were anesthetized with isoflurane and nasal washes were collected with 400 µl of PBS on endpoint days (2, 4, and 7 dpi). Hamsters were euthanized by CO_2_ for organ collection. Nasal wash and lung were collected to measure viral titer and RNA. Left lungs were collected for histopathology.

### Histology

Left lung lobes were harvested from hamsters and fixed in 10% buffered formalin solution for at least 7 days. Fixed tissue was then embedded in paraffin, cut into 5 µM sections, and stained with hematoxylin and eosin (H&E) on a SAKURA VIP6 processor by the University of Texas Medical Branch Surgical Pathology Laboratory.

### Immunohistochemistry

Fixed and paraffin-embedded left lung lobes from hamsters were cut into 5 µM sections and mounted onto slides by the University of Texas Medical Branch Surgical Pathology Laboratory. Paraffin-embedded sections were warmed at 56°C for 10 min, deparaffinized with xylene (3x 5-min washes) and graded ethanol (3x 100% 5-min washes, 1x 95% 5-min wash), and rehydrated in distilled water. After rehydration, antigen retrieval was performed by steaming slides in antigen retrieval solution (10 mM sodium citrate, 0.05% Tween-20, pH 6) for 40 min (boil antigen retrieval solution in microwave, add slides to boiling solution, and incubate in steamer). After cooling and rinsing in distilled water, endogenous peroxidases were quenched by incubating slides in TBS with 0.3% H_2_O_2_ for 15 min followed by 2x 5-min washes in 0.05% TBST. Sections were blocked with 10% normal goat serum in BSA diluent (1% BSA in 0.05% TBST) for 30 min at room temperature. Sections were incubated with primary anti-N antibody (Sino #40143-R001) at 1:1000 in BSA diluent overnight at 4°C. Following overnight primary antibody incubation, sections were washed 3x for 5 min in TBST. Sections were incubated in secondary HRP-conjugated anti-rabbit antibody (Cell Signaling Technology #7074) at 1:200 in BSA diluent for 1 hour at room temperature. Following secondary antibody incubation, sections were washed 3x for 5 min in TBST. To visualize antigen, sections were incubated in ImmPACT NovaRED (Vector Laboratories #SK-4805) for 3 min at room temperature before rinsed with TBST to stop the reaction followed by 1x 5-min wash in distilled water. Sections were incubated in hematoxylin for 5 min at room temperature to counterstain before rinsing in water to stop the reaction. Sections were dehydrated by incubating in the previous xylene and graded ethanol baths in reverse order before mounted with coverslips.

### Immunofluorescence

Hela or HeLa-FRT UBAP2L KO or parental cells were seeded in 6-well dishes with coverslips at 25% confluency. Cells were transfected the day after with 250 ng DNA and 1 μL or jet OPTIMUS (Polyplus) reagent overnight in DMEM media supplemented with FBS (10%) (HyClone) and PenStrep (1%) (Thermo Scientific). Media was changed to media containing 0,5 μM sodium arsenite for 30 minutes to induce stress granules formation. After washing with PBS cells were fixed for 20 minutes with 4% formaldehyde in PBS. Cells were premetallized for 10 minutes with PBS 0,5% Triton-100. Following three 5-minute washes with PBS-T (0,05% Tween), 25 mM Glycine was incubated overnight at 4°C. Coverslips were blocked in TBS-T (0,05% Tween) 3% BSA for 45 minutes at room temperature. Primary antibodies (anti-G3BP1 mouse abcam #ab56574 or anti-FXR1 mouse clone 6BG10 Milipore #05-1529) were incubated at 1:400 dilution in 3% BSA TBS-T(0,05% Tween) overnight at 4°C. Following 3three 5-minute washes with TBS-T (0,05% Tween), coverslips were incubated with secondary antibodies for 1h at room temperature. Coverslips were mounted in MOWIOL mounting solution (Calbiochem #475904) and imaged on a Delta-Vision Elite microscope (DeltaVision) with 60x oil objective. Data was analyzed in Fiji and plotted with a Prism 9 GraphPad software.

### Live cell imaging

HeLa cells were seeded at 25% confluency 6-well dishes and transfected on the following day with 200 ng DNA and 1μL JetOptimus reagent overnight in 3mL media. Media was changed on the next day and cells seeded in 8-well ibidi dishes at 40% confluency. Pictures were taken every 5-10 minutes in 2 z-stacks on a Delta-Vision Elite microscope (DeltaVision) with 60x oil objective. Cells were filmed 48 hours post transfection for 10 minutes and sodium arsenite was added to a final concentration of 0,5 μM. Cells fere filmed for additional hour to follow the formation of stress granules. Data was analyzed in Fiji and plotted with a Prism 9 GraphPad software.

### Immunoprecipitations

Cells (HeLa or HEK293) were seeded at 25% confluency in 15 cm^3^ dishes with 2 μg DNA and 2μL Jet Optimus reagent overnight in 15mL media. Media was changed on the following day and cells were collected after 48 or 24 hours post transfection. Cell pellets from each 15 cm^3^ dish were lyzed in 350 μL lysis buffer (100mM NaCl, 50mM Tris pH 7,4, 0,1% NP40, 0,2% Triton-100, 1mM DTT) supplemented with protease (complete mini EDTA free, Roche) and phosphatase inhibitor tablets (Roche). Lysate was sonicated at 4°C for 10 cycles of 30s ON, 30s OFF using a Bioruptor sonicator. Following sonication lysates were cleared for 45 min at 20000g. Supernatants were collected and concentrations were measured. Lysates were incubated with 10 μL pre-equilibrated GFP-trap beads for 1h at 4°C on a rotor-wheel. Beads were washed 3 times with 800 μL wash buffer (150mM NaCl, 50mM Tris pH 7,4, 0,05% NP40, 5% Glycerol, 1mM DTT). If immunoprecipitations were prepared for Mass-spectrometry analysis, one additional wash with basic wash buffer (100mM NaCl, 50mM Tris pH 7,4, 5% Glycerol) was made. If immunoprecipitations were analyzed by SDS-PAGE and western blot, 25 μL 2xLSB (Thermo Fisher Scientific) was used to elute the samples. Samples were then boiled and separated by SDS-PAGE.

### Size-exclusion chromatography

Recombinant proteins (GST-NSP3 1-181 and His-FXR1 215-360) were ran on a Supperose 200 column (Cytiva) and an AKTA system. For analyzing direct protein-protein interactions, proteins were pre-mixed for 30 min on ice and then following a 30s spin at 20000g on a table-top centrifuge they were ran on the column. 500 μL fractions were collected and peak fractions were analyzed by SDS-PAGE.

### Structural modelling

Structures of complexes between human FXR1 and NSP3 or UBA2PL were predicted with Alphafold multimer [40, 51, 52] based on full-length amino acid sequences for human FXR1 (UniProt entry A0A0F7L1S3) and human UBA2PL (Uniprot entry F8W726) and SarsCov2 NSP3 residues 103-161 (Uniprot entry P0DTD1).

Phosphorylations of Serine and Threonine residues were modelled and locally geometry-refined in Coot [53]. All structural models/PDBs and their pLDDT scores were visualized in PyMOL (Schrodinger, LLC).

#### Isothermal Titration Calorimetry (ITC)

Peptides were purchased from Peptide 2.0 Inc (Chantilly. VA, USA). The purity obtained in the synthesis was 95 – 98 % as determined by high performance liquid chromatography (HPLC) and subsequent analysis by mass spectrometry. Prior to ITC experiments both the proteins and the peptides were extensively dialyzed against 50 mM sodium phosphate pH 7.5, 150 mM NaCl, 0.5 mM TCEP. All ITC experiments were performed on an Auto-iTC200 instrument (Microcal, Malvern Instruments Ltd.) at 25 °C. Both peptide and protein concentrations were determined using a spectrometer by measuring the absorbance at 280 nm and applying values for the extinction coefficients computed from the corresponding amino acid sequences by the ProtParam program (http://web.expasy.org/protparam/). FXR1 constructs (FXR1*^212-289^*, FXR1*^215-360^*) and NCAP_SARS2*^1-419^*at approximately 300 μM or 100 μM (FXR1*^1-122^*) concentration were loaded into the syringe and titrated into the calorimetric cell containing NSP3 1-181 at ∼ 20 μM or ∼ 10 μM, respectively. NSP3 128-148 peptide (and variants) and UBAP2L 243-270 (and variants) at approximately 300 μM were loaded into the syringe and titrated into the calorimetric cell containing FXR1*^215-360^* at ∼ 20 μM. For competition experiments, NSP3 1-181 at approximately 300 μM concentration was loaded into the syringe and titrated into the calorimetric cell containing either FXR1*^215-360^* or FXR1*^215-360^* saturated with UBAP2L 243-270 at ∼ 20 μM FXR1*^215-360^* concentration. The reference cell was filled with distilled water. In all assays, the titration sequence consisted of a single 0.4 μl injection followed by 19 injections, 2 μl each, with 150 s spacing between injections to ensure that the thermal power returns to the baseline before the next injection. The stirring speed was 750 rpm. Control experiments with the FXR1, NCAP_SARS2 constructs or the NSP3 and UBAP2L peptides injected in the sample cell filled with buffer were carried out under the same experimental conditions. These control experiments showed heats of dilution negligible in all cases. The heats per injection normalized per mole of injectant *versus* the molar ratio [titrant in syringe]/[titrand in calorimetric cell] were fitted to a single-site model. RNA sequence for RNA used in S3A: rGrGrArUrCrArUrUrUrUrGrUrUrGrGrArCrUrCrArArUrUrUrCrArArCrUrCrUrArArCrUrU rUrArArCrUrUrUrGrCrArUrUrGrGrUrUrGrGrArCrArCrCrU. Data were analysed with MicroCal PEAQ-ITC (version 1.1.0.1262) analysis software (Malvern Instruments Ltd.).

#### Protein production and purification

The NSP3 and FXR1 fragments were expressed in the *E. coli.* strain BL21(DE3) overnight at 18 degrees. Cells were harvested and resuspended in 50 mM NaP pH=7,5; 300 mM NaCl; 10% glycerol; 0,5 mM TCEP; 1XComplete EDTA-free tablets (Roche) (and 10 mM imidazole for His-tag purifications) and lysed with high-pressure-homogonizer (Avestin) and cell extract clarified by centrifugation. The clarified lysate was loaded onto a His-tag or GST-tag affinity column and following washing with resuspension buffer the proteins were eluted with either an imidazole gradient or glutathione containing buffer. Peak fractions were pooled and further purified on a size-exclusion chromatography column pre-equilibrated with 50 mM NaP pH=7,5; 150 mM NaCl; 10% glycerol; 0,5 mM TCEP.

### Affinity purification and mass spectrometry (AP-MS)

Partial on-bead digestion was used for peptide elution from GFP-Trap Agarose (Chromotek). Briefly, 100 μl of elution buffer (2 M urea; 2 mM DTT; 20 μg/ml trypsin; and 50 mM Tris, pH 7.5) was added and incubated at 37°C for 30 min. Samples were alkylated with 25 mM CAA and digested overnight at room temperature before addition of 1% trifluoroacetic acid (TFA) to stop digestion. Peptides were desalted and purified with styrene–divinylbenzene reversed-phase sulfonate (SDB-RPS) StageTips. Briefly, two layers of SDB-RPS were prepared with 100 μl wash buffer (0.2% TFA in H2O). Peptides were loaded on top and centrifuged for 5 min at 500 g, and washed with 150 μl wash buffer. Finally, peptides were eluted with 50 μl elution buffer (80% ACN and 1% ammonia) and vacuum-dried. Dried peptides were dissolved in 2% acetonitrile (ACN) and 0.1% TFA in water and stored at −20°C.

### LC-MS analysis

Liquid chromatography mass spectrometry (LC-MS) analysis was performed with an EASY-nLC-1200 system (Thermo Fisher Scientific) connected to a trapped ion mobility spectrometry quadrupole time-of-flight mass spectrometer (timsTOF Pro, Bruker Daltonik GmbH, Germany) with a nano-electrospray ion source (Captive spray, Bruker Daltonik GmbH). Peptides were loaded on a 50 cm in-house packed HPLC-column (75µm inner diameter packed with 1.9µm ReproSilPur C18-AQ silica beads, Dr. Maisch GmbH, Germany). Peptides were separated using a linear gradient from 5-30% buffer B (0.1% formic acid, 80% ACN in LC-MS grade H2O) in 43 min followed by an increase to 60% buffer B for 7 min, then to 95% buffer B for 5min and back to 5% buffer B in the final 5min at 300nl/min. Buffer A consisted of 0.1% formic acid in LC-MS grade H2O. The total gradient length was 60 min. We used an in-house made column oven to keep the column temperature constant at 60 °C.

Mass spectrometric analysis was performed essentially as described in Brunner et al. [54] in data-dependent (ddaPASEF) mode. For ddaPASEF, 1 MS1 survey TIMS-MS and 10 PASEF MS/MS scans were acquired per acquisition cycle. Ion accumulation and ramp time in the dual TIMS analyzer was set to 100 ms each and we analyzed the ion mobility range from 1/K0 = 1.6 Vs cm-2 to 0.6 Vs cm-2. Precursor ions for MS/MS analysis were isolated with a 2 Th window for m/z < 700 and 3 Th for m/z >700 in a total m/z range of 100-1.700 by synchronizing quadrupole switching events with the precursor elution profile from the TIMS device. The collision energy was lowered linearly as a function of increasing mobility starting from 59 eV at 1/K0 = 1.6 VS cm-2 to 20 eV at 1/K0 = 0.6 Vs cm-2. Singly charged precursor ions were excluded with a polygon filter (otof control, Bruker Daltonik GmbH). Precursors for MS/MS were picked at an intensity threshold of 1.000 arbitrary units (a.u.) and resequenced until reaching a ‘target value’ of 20.000 a.u taking into account a dynamic exclusion of 40 s elution.

### Data analysis of proteomic raw files

Mass spectrometric raw files acquired in ddaPASEF mode were analyzed with MaxQuant (version 1.6.7.0) [55, 56]. The Uniprot database (2019 release, UP000005640_9606) was searched with a peptide spectral match (PSM) and protein level FDR of 1%. A minimum of seven amino acids was required including N-terminal acetylation and methionine oxidation as variable modifications and cysteine carbamidomethylation as fixed modification. Enzyme specificity was set to trypsin with a maximum of two allowed missed cleavages. First and main search mass tolerance was set to 70 ppm and 20 ppm, respectively. Peptide identifications by MS/MS were transferred by matching four-dimensional isotope patterns between the runs (MBR) with a 0.7-min retention-time match window and a 0.05 1/K0 ion mobility window. Label-free quantification was performed with the MaxLFQ algorithm [57] and a minimum ratio count of two.

### Bioinformatic analysis

Proteomics data analysis was performed with Perseus [58] and within the R environment (https://www.r-project.org/). MaxQuant output tables were filtered for ‘Reverse’, ‘Only identified by site modification’, and ‘Potential contaminants’ before data analysis. Missing values were imputed after stringent data filtering and based on a normal distribution (width = 0.3; downshift = 1.8) prior to statistical testing. For pairwise proteomic comparisons (two-sided unpaired t-test), we applied a permutation-based FDR of 5% to correct for multiple hypothesis testing including an *s_0_* value [59] of 0.1.

## Figure legends

**Supplemental Figure 1.**
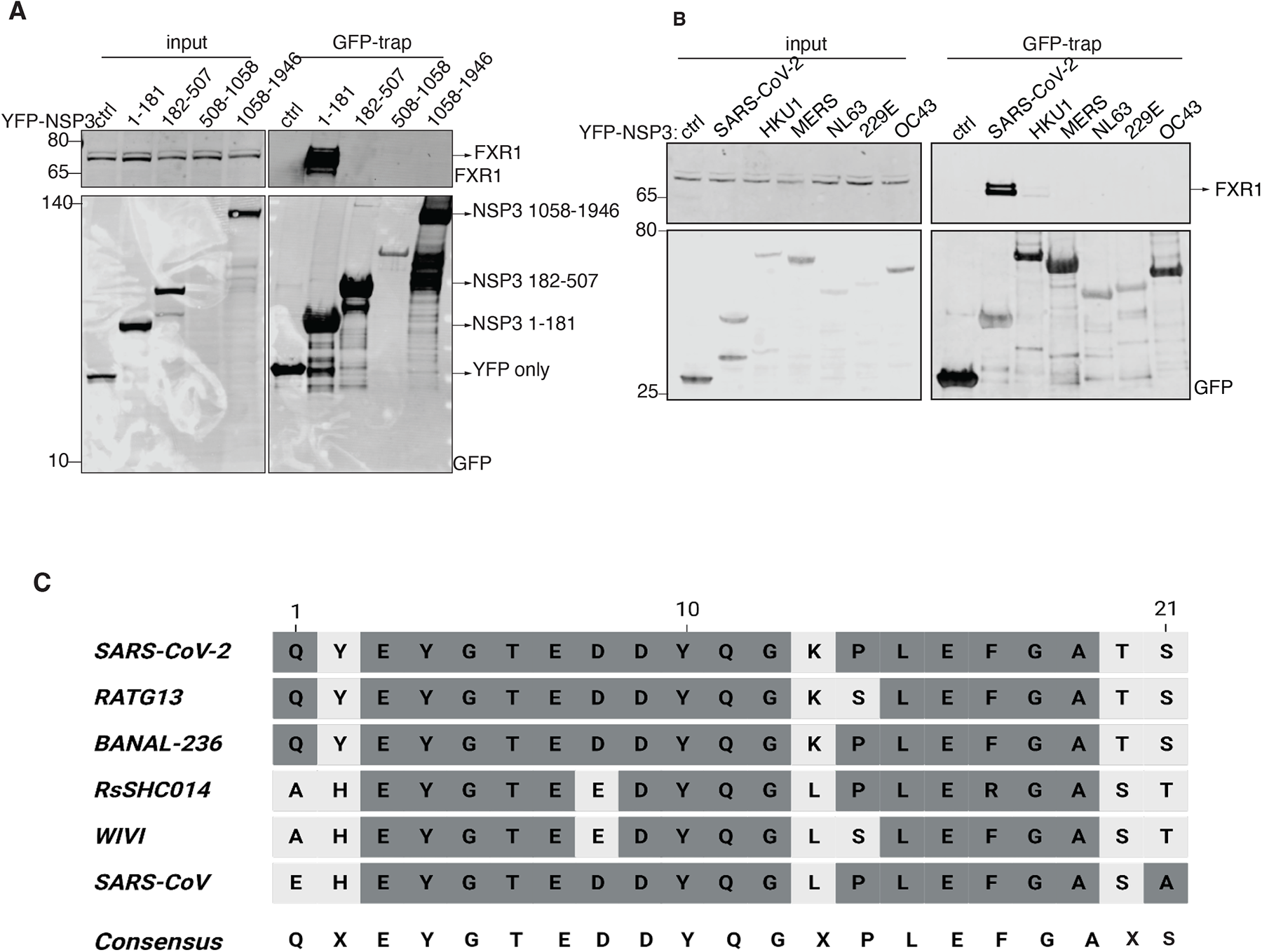
Analysis of FMRP-NSP3 interaction. A) The indicated NSP3 fragments fused to YFP was expressed and purified from HeLa cells and binding to FXR1 monitored. Representative of at least 2 experiments. B) A panel of NSP3 N-terminal fragments from different coronaviruses were expressed and purified from HeLa cells and binding to FXR1 determined by western-blotting. Representative of 2 independent experiments. C) Alignment of the NSP3 sequence binding to FMRPs from different coronaviruses.

**Supplemental Figure 2.**
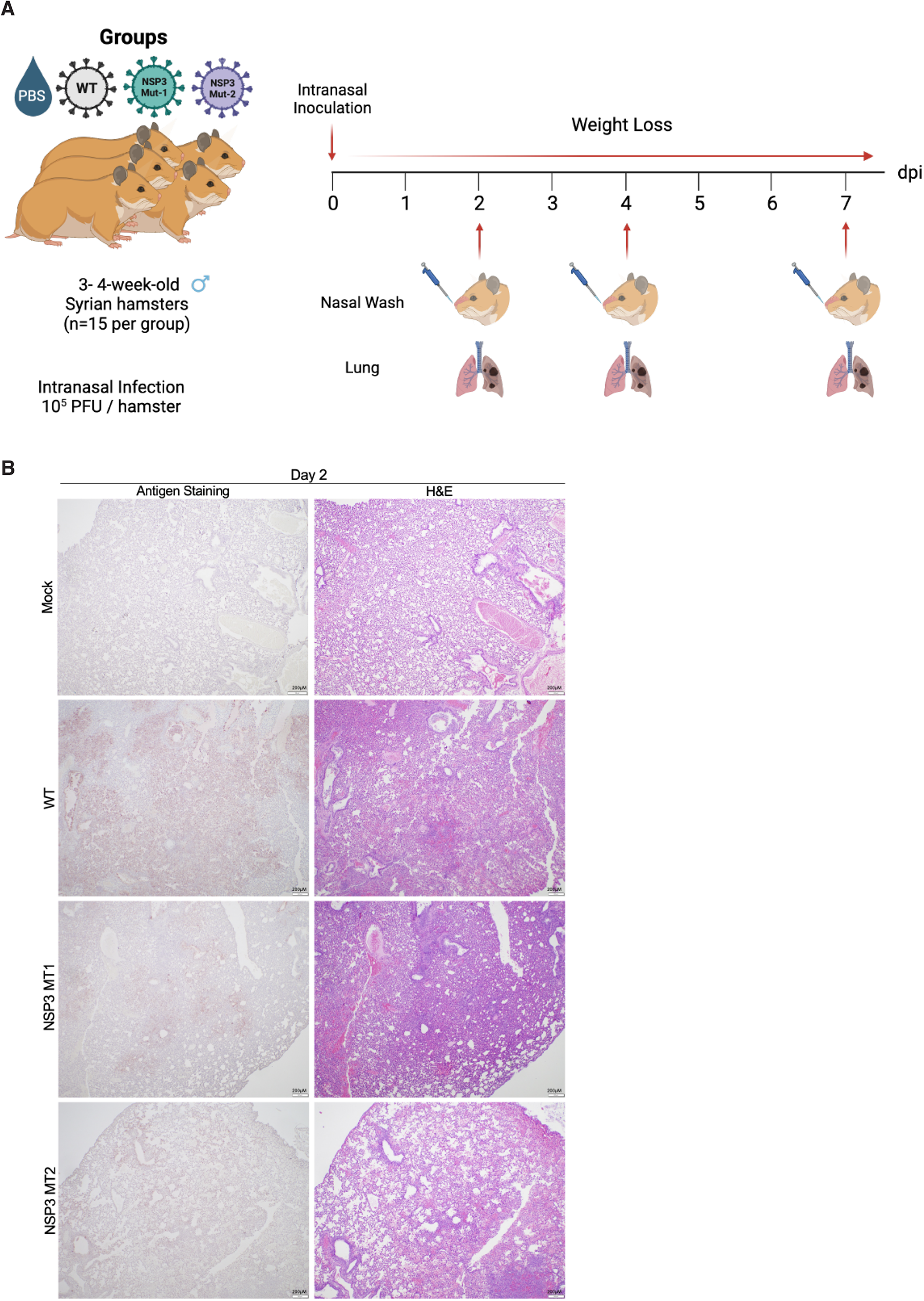
Histopathology of hamster infected with WT or NSP3 Mutants. A) Schematic of in vivo experiment (generated with BioRender). B) H&E and viral antigen (nucleocapsid) immunohistochemical staining of lung of hamsters infected mock (PBS) or with 10^5^ pfu of WT, NSP3 Mut1, or NSP3 Mut 2 SARS-CoV-2 at 2 days post infection. WT infection shows extensive viral infection and damage; both NSP3 mutants have focal disease and less damage. No damage observed in mock infected samples.

**Supplemental Figure 3.**
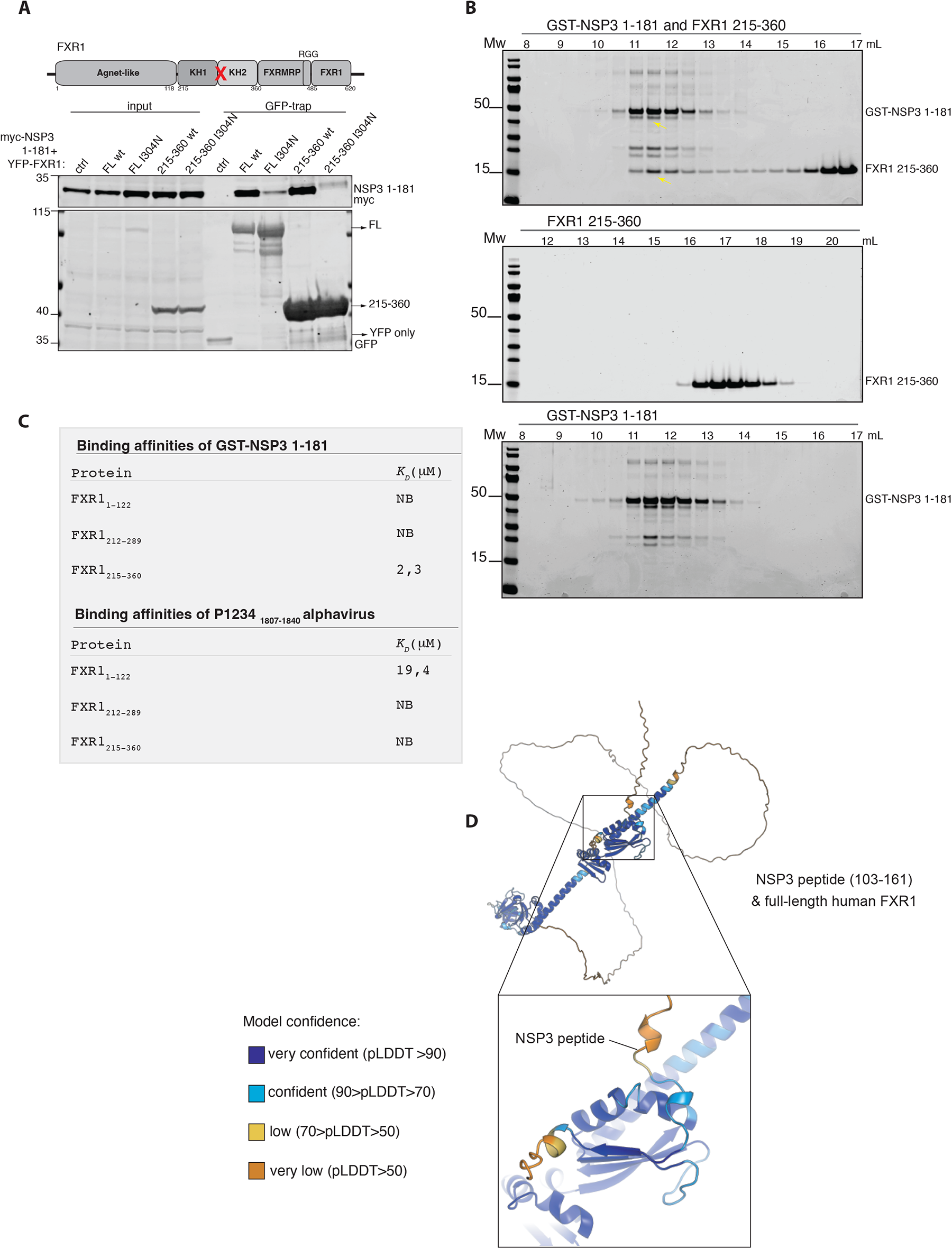
A direct interaction between NSP3 and FXR1. A) The indicated FXR1 YFP constructs were co-expressed with myc-NSP3 1-181 in HeLa cells and affinity purified using YFP affinity beads. The binding to NSP3 was monitored by probing for myc. Representative of 2 independent experiments. B) Size exclusion chromatography of GST-NSP3 WT 1-181, FXR1 215-360 either alone or in combination. The elution volume is indicated on top and coomasie stained gels of fractions shown. Representative of 2 independent experiments. C) Table of ITC values obtained for the indicated FXR1 fragments binding to GST-NSP3 1-181 or the FXR1 binding peptide from old alphaviruses. D) Confidence plots of AlphaFold model of NSP3 peptide binding to full length FXR1.

**Supplemental Figure 4.**
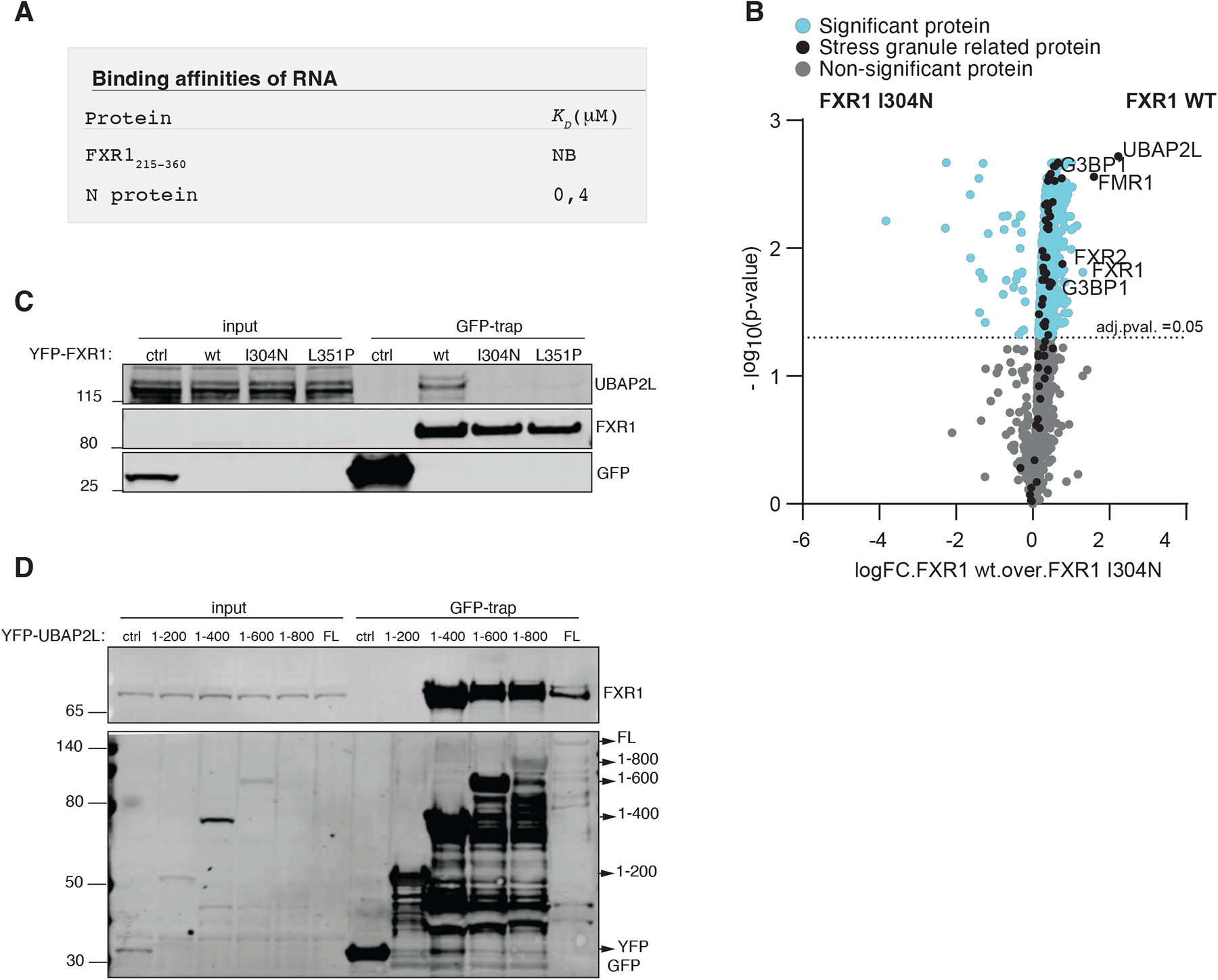
Interaction of FXR1 to UBAP2L. A) ITC measurements of a reported RNA binding to FXR1. Binding to FXR1 215-360 was monitored and as a control the SARS-CoV-2 N protein. B) Mass spectrometry analysis of the interactomes of affinity purified YFP-tagged FXR1 WT and FXR1 I304N. Proteins specifically binding to FXR1 WT indicated in the volcano plot. Data from 4 technical repeats. C) The indicated YFP-tagged FXR1 proteins were expressed and purified from HeLa cells and binding to UBAP2L determined by western-blot. D) A panel of YFP-UBAP2L constructs were expressed and purified from HeLa cells and binding to FXR1 determined. Representative of 2 independent experiments.

**Supplemental Figure 5.**
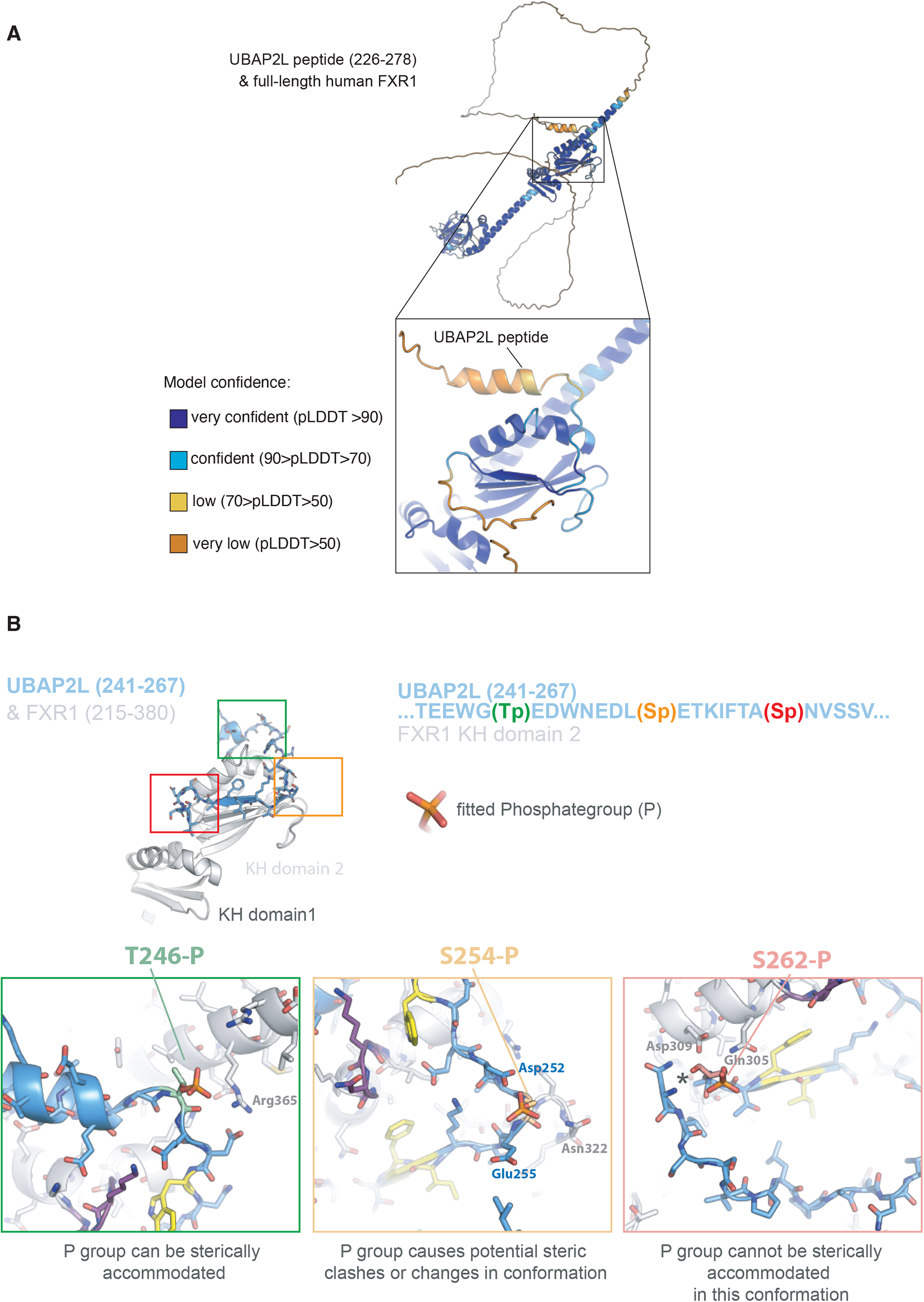
Model of interaction of FXR1 to UBAP2L. A) Confidence plots of AlphaFold model of UBAP2L peptide 226-278 binding to full length FXR1. B) Reported phosphorylation sites in UBAP2L were fitted into the AlphaFold of the FXR1-UBAP2L complex.

**Supplemental Figure 6.**
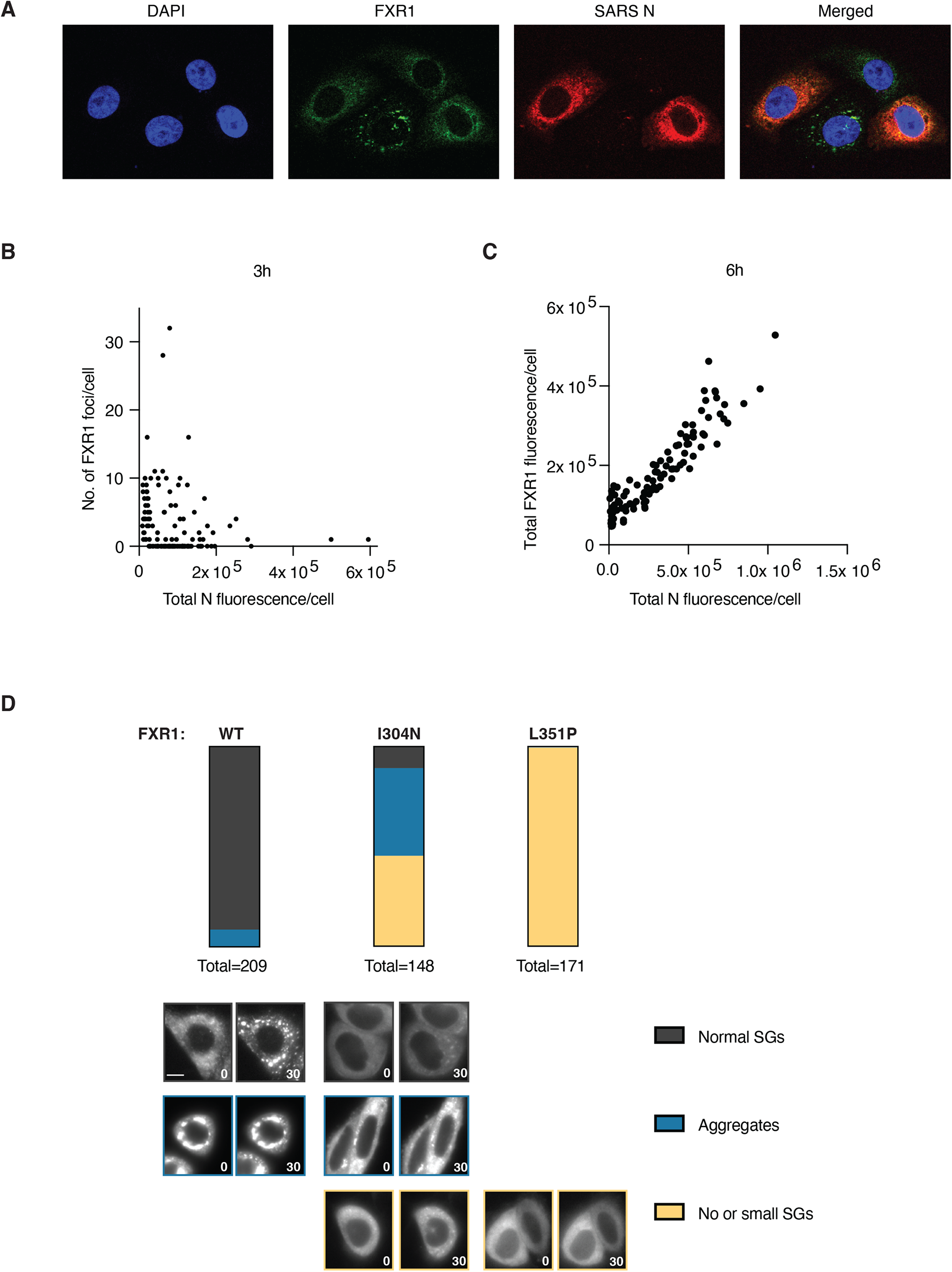
Analysis of FXR1 localisation to stress granules. A) Vero E6 cells infected with SARS-CoV-2 were fixed and stained for FXR1 and the viral N protein. Representative images shown. B) The number of FXR1 foci in infected cells was determined and correlated with total level of N protein. C) The total level of FXR1 was determined in infected cells and plotted against total levels of N. D) YFP-tagged FXR1 proteins were expressed in HeLa cells and filmed by live cell microscopy. Stress granule formation was induced by arsenite and 30 minutes after addition the localization and morphology of FXR1 foci was monitored. Phenotypes are plotted as percentage. Scores of two individual experiments are shown. The total number of cells analyzed per condition are indicated. Representative images are shown with a scale bar of 10 μM indicated in lower left corner.

**Supplemental Figure 7.**
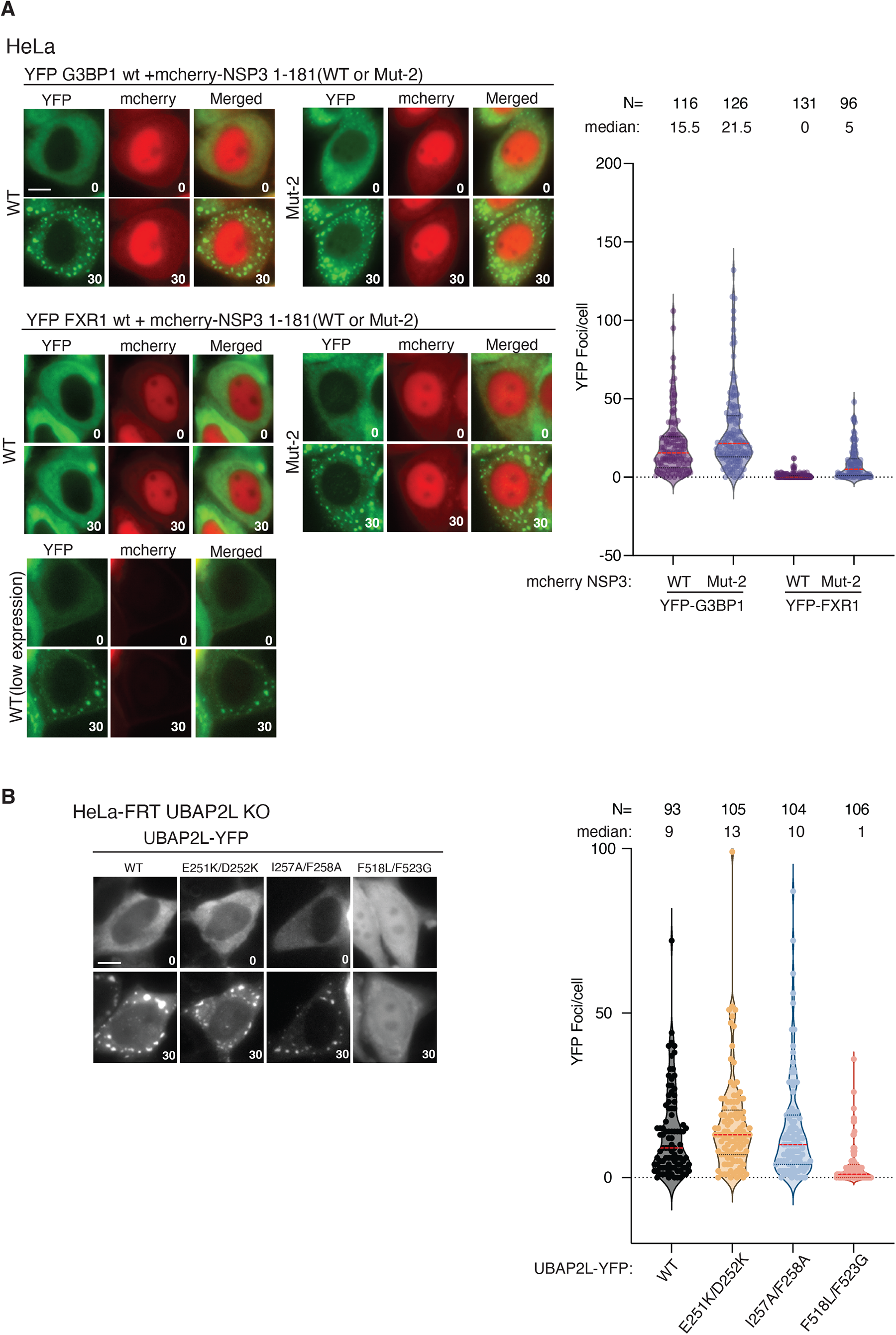
Analysis of stress granule components by live cell microscopy. A) The indicated YFP tagged stress granule proteins were expressed in HeLa cells in the presence of mCherry tagged NSP3 constructs and monitored by live cell microscopy. Stress granule formation was induced by arsenite and number of stress granules determined after 30 min. A pool of 3 experiments is shown in the graph. B) The indicated UBAP2L-YFP constructs were expressed in HeLa cells and monitored by live cell microscopy. Arsenite was added and localization determined after 30 minutes. The combined data from 2 experiments is shown in the graphs. In A-B representative stills from the immunofluorescence is shown with a scale bar of 10 μM indicated in lower left corner. The median is indicated with red line. The median and number of cells (N) analyzed per condition is indicated above the plot.

