## Supplementary material for "SARS-CoV-2 hijacks fragile X mental retardation proteins for efficient infection": Table 2

***Table 2.*** Affinities and thermodynamic values of NSP3 1-181 binding events inferred from ITC measurements performed at 25 °C. Gibbs free energy (ΔG), enthalpy (ΔH), entropy (-TΔS), equilibrium dissociation constant ($K_D$) and reaction stoichiometry (n) are shown. The protein-protein interaction affinity is defined by the Gibbs energy for binding $\Delta G = -RT \ln K_A = RT \ln K_D$.

| | **ΔG (*kcal mol⁻¹*)** | **ΔH (*kcal mol⁻¹*)** | **-TΔS (*kcal mol⁻¹*)** | **$K_D$ (*μM*)** | ***n*** |
|---|---|---|---|---|---|
| NSP3 1-181/FXR1$^{1\text{-}122}$ | | | *No binding* | | |
| NSP3 1-181/FXR1$^{212\text{-}289}$ | | | *No binding* | | |
| NSP3 1-181/FXR1$^{215\text{-}360}$ | -7.70 | -25.9 ± 0.38 | 18.2 | 2.26 ± 0.13 | 0.90 ± 0.01 |
| NSP3 1-181mut1/FXR1$^{215\text{-}360}$ | | | *No binding* | | |
| NSP3 1-181/FXR1$^{215\text{-}360}$ I304N | | | *No binding* | | |

***Table 2 continued.*** Affinities and thermodynamic values of FXR1$^{215\text{-}360}$ binding events inferred from ITC measurements performed at 25 °C. Gibbs free energy (ΔG), enthalpy (ΔH), entropy (-TΔS), equilibrium dissociation constant ($K_D$) and reaction stoichiometry (n) are shown. The protein-protein interaction affinity is defined by the Gibbs energy for binding $\Delta G = -RT \ln K_A = RT \ln K_D$. [a]The stoichiometry (n) was fixed to 1 when not explicitly stated.

| FXR1$^{215\text{-}360}$ complex | **ΔG (*kcal mol*$^{-1}$)** | **ΔH (*kcal mol*$^{-1}$)** | **-TΔS (*kcal mol*$^{-1}$)** | **$K_D$ (*μM*)** | ***n*** |
|---|---|---|---|---|---|
| FXR1$^{215\text{-}360}$/NSP3 WT peptide | -7.86 | -24.20 ± 0.12 | 16.3 | 1.74 ± 0.04 | 0.95 ± 0.01 |
| FXR1$^{215\text{-}360}$/NSP3 M1 | -5.49 | -18.40 ± 8.67 | 13.0 | 95.1 ± 43.2 | |
| FXR1$^{215\text{-}360}$/NSP3 M2 | | | *No binding* | | |
| FXR1$^{215\text{-}360}$/NSP3 M3 | | | *No binding* | | |
| FXR1$^{215\text{-}360}$/NSP3 M4 | | | *No binding* | | |

***Table 2 continued.*** Affinities and thermodynamic values of NSP3 alphavirus binding events inferred from ITC measurements performed at 25 °C. Gibbs free energy (ΔG), enthalpy (ΔH), entropy (-TΔS), equilibrium dissociation constant ($K_D$) and reaction stoichiometry (n) are shown. The protein-protein interaction affinity is defined by the Gibbs energy for binding $\Delta G = -RT \ln K_A = RT \ln K_D$. [a]The stoichiometry (n) was fixed to 1 when not explicitly stated.

| | **ΔG (*kcal mol$^{-1}$*)** | **ΔH (*kcal mol$^{-1}$*)** | **-TΔS (*kcal mol$^{-1}$*)** | **$K_D$ (*μM*)** | ***[a]n*** |
|---|---|---|---|---|---|
| NSP3 peptide alphavirus/FXR1$^{1-122}$ | -6.43 | -6.11 ± 0.65 | -0.32 | 19.4 ± 3.16 | |
| NSP3 peptide alphavirus/FXR1$^{212-289}$ | | | *No binding* | | |
| NSP3 peptide alphavirus/FXR1$^{215-360}$ | | | *No binding* | | |

***Table 2 continued.*** Affinities and thermodynamic values of FXR1$^{215\text{-}360}$ binding events inferred from ITC measurements performed at 25 °C. Gibbs free energy (ΔG), enthalpy (ΔH), entropy (-TΔS), equilibrium dissociation constant ($K_D$) and reaction stoichiometry (n) are shown. The protein-protein interaction affinity is defined by the Gibbs energy for binding $\Delta G = -RT \ln K_A = RT \ln K_D$. [a]The stoichiometry (n) was fixed to 1 when not explicitly stated.

| FXR1$^{215\text{-}360}$ complex | **ΔG (*kcal mol$^{-1}$*)** | **ΔH (*kcal mol$^{-1}$*)** | **-TΔS (*kcal mol$^{-1}$*)** | **$K_D$ (*µM*)** | ***n*** |
|---|---|---|---|---|---|
| FXR1$^{215\text{-}360}$/UBAP2L$^{263\text{-}290}$ | -6.92 | -13.7 ± 0.54 | 6.79 | 8.53 ± 0.69 | 0.96 ± 0.01 |
| FXR1$^{215\text{-}360}$/UBAP2L$^{263\text{-}290}$ *pS274* | -6.27 | -8.91 ± 2.38 | 2.65 | 25.5 ± 7.34 | 0.92 ± 0.01 |
| FXR1$^{215\text{-}360}$/UBAP2L$^{263\text{-}290}$ *pT266* | -7.42 | -12.0 ± 0.26 | 4.57 | 3.66 ± 0.24 | 0.90 ± 0.01 |
| FXR1$^{215\text{-}360}$/UBAP2L$^{263\text{-}290}$ *pS282* | | | *No binding* | | |

***Table 2 continued*****.** Affinities and thermodynamic values of NF1/N protein from SARS-CoV2 and FXR1 215-360 binding inferred from ITC measurements performed at 25 °C. Gibbs free energy (ΔG), enthalpy (ΔH), entropy (-TΔS), equilibrium dissociation constant ($K_D$) and reaction stoichiometry (n) are shown. The protein-protein interaction affinity is defined by the Gibbs energy for binding $\Delta G = -RT \ln K_A = RT \ln K_D$. [a]The stoichiometry (n) was fixed to 1 when not explicitly stated.

| FXR1$^{215-360}$ complex | **ΔG (*kcal mol*$^{-1}$)** | **ΔH (*kcal mol*$^{-1}$)** | **-TΔS (*kcal mol*$^{-1}$)** | **$K_D$ (*μM*)** | ***n*** |
|---|---|---|---|---|---|
| N SARS-CoV-2/NF1 RNA | -8.76 | -30.1 ± 0.68 | 21.3 | 0.38 ± 0.04 | 0.82 ± 0.01 |
| FXR1$^{215-360}$/NF1 RNA | | | *No binding* | | |

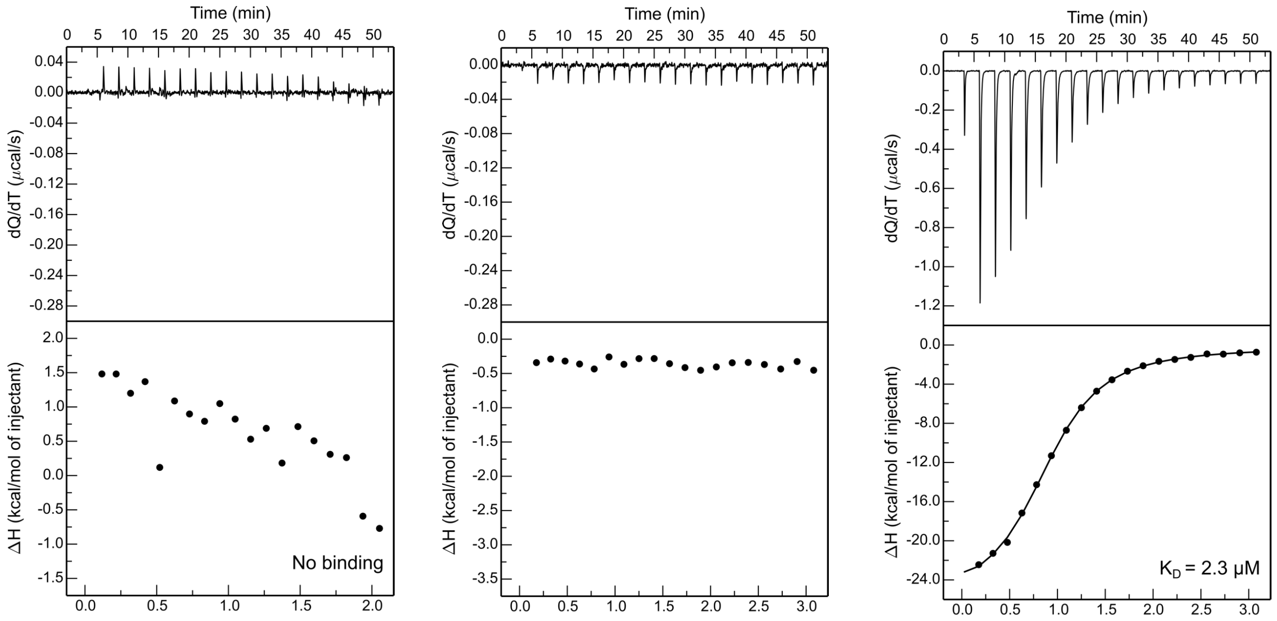


FXR1 1-122/GST-NSP3 1-181

FXR1 212-289/GST-NSP3 1-181

FXR1 215-360/GST-NSP3 1-181

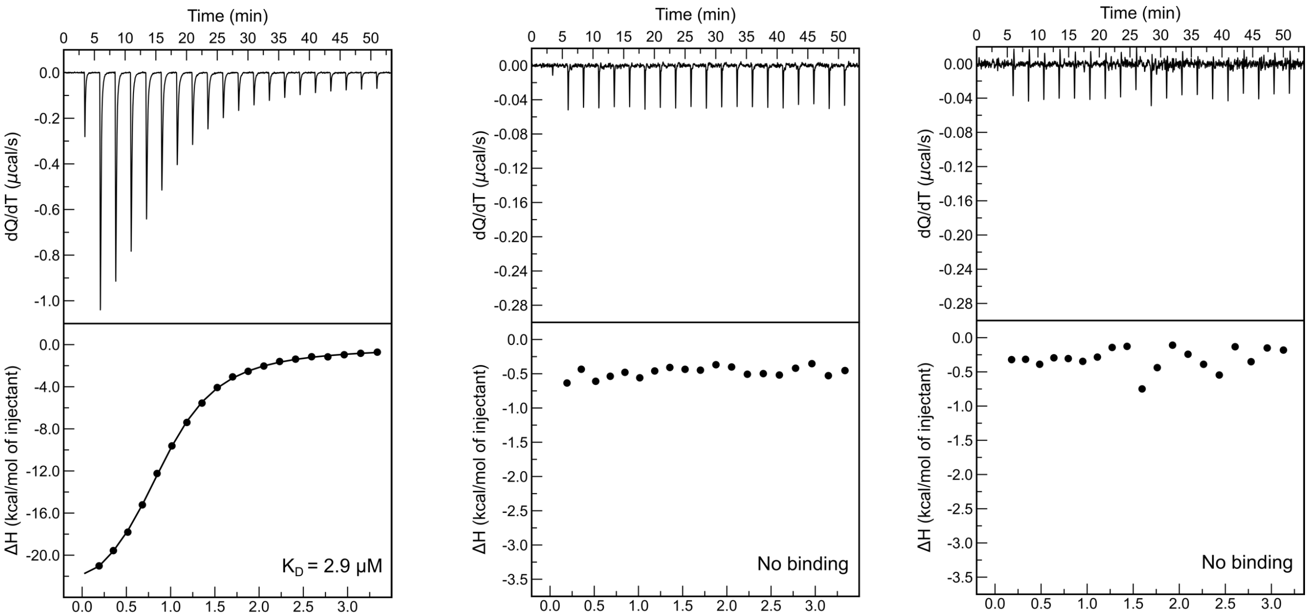


FXR1 215-360/His-NSP3 1-181 WT

FXR1 215-360/His-NSP3 1-181 Mut1

FXR1 215-360 I304N/His-NSP3 1-181

***Table 2 continued***

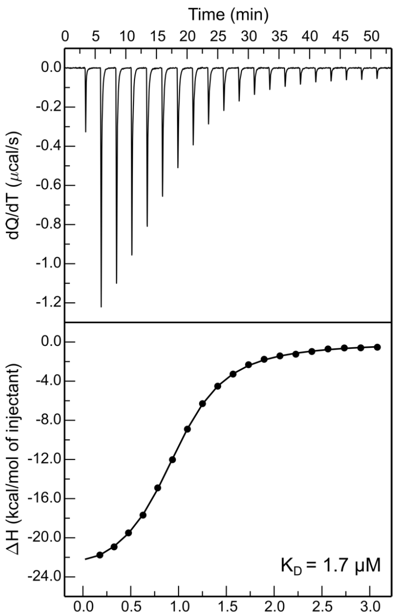


FXR1 215-360/NSP3 peptide WT

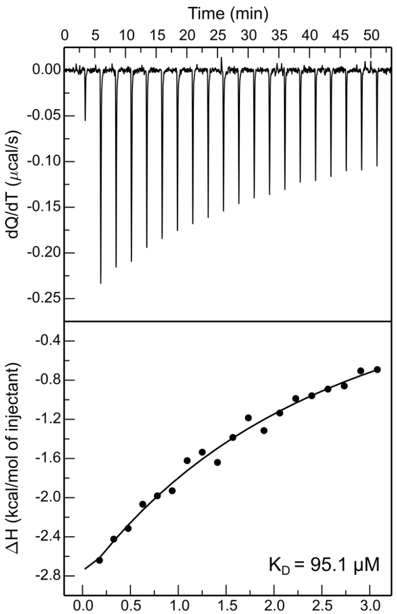


FXR1 215-360/NSP3 peptide M1

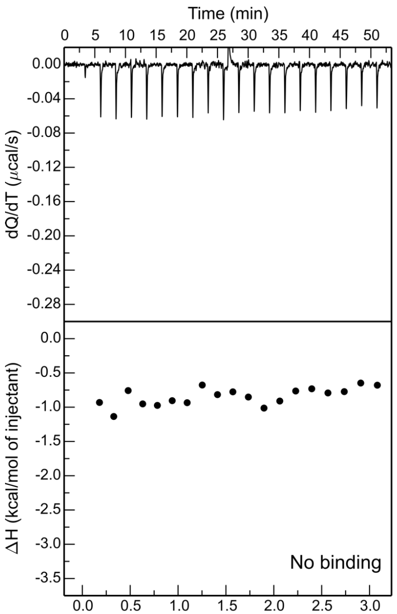


FXR1 215-360/NSP3 peptide M2

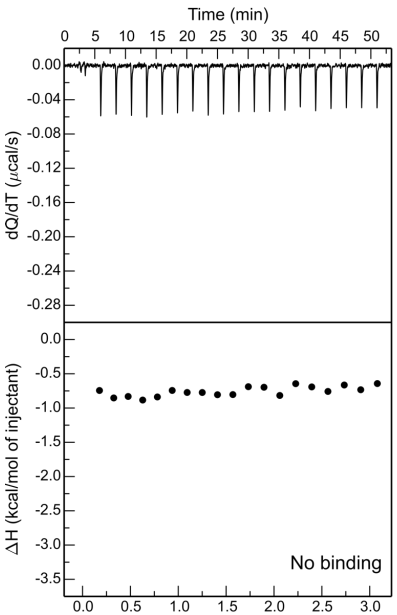


FXR1 215-360/NSP3 peptide M3

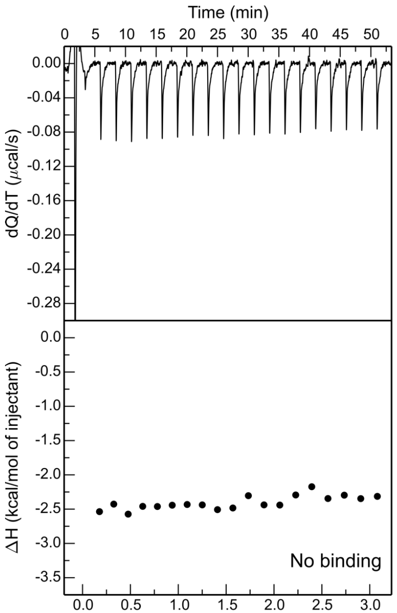


FXR1 215-360/NSP3 peptide M4

***Table 2 continued***

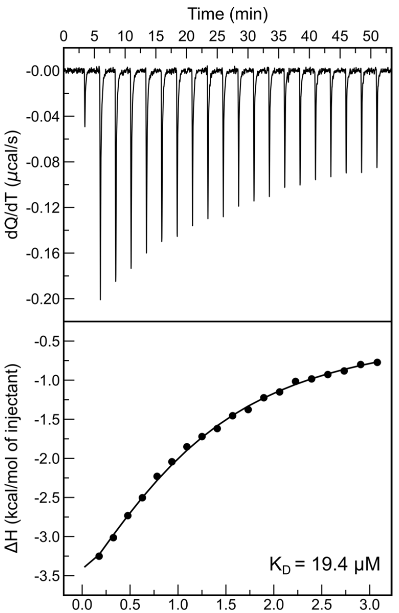


FXR1 1-122/NSP3 peptide alphavirus

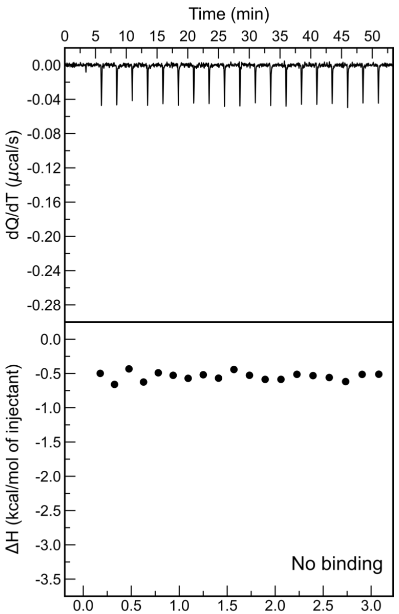


FXR1 212-289/NSP3 peptide alphavirus

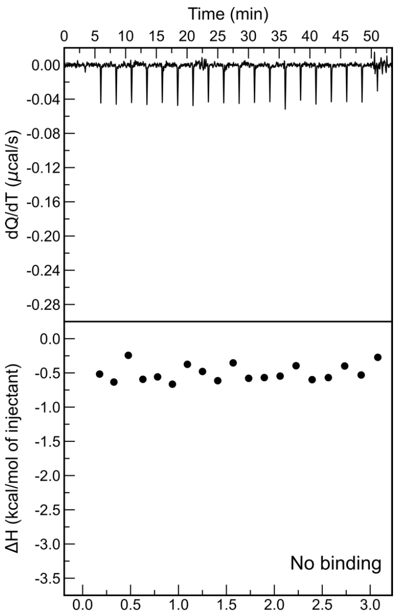


FXR1 215-360/NSP3 peptide alphavirus

***Table 2 continued***

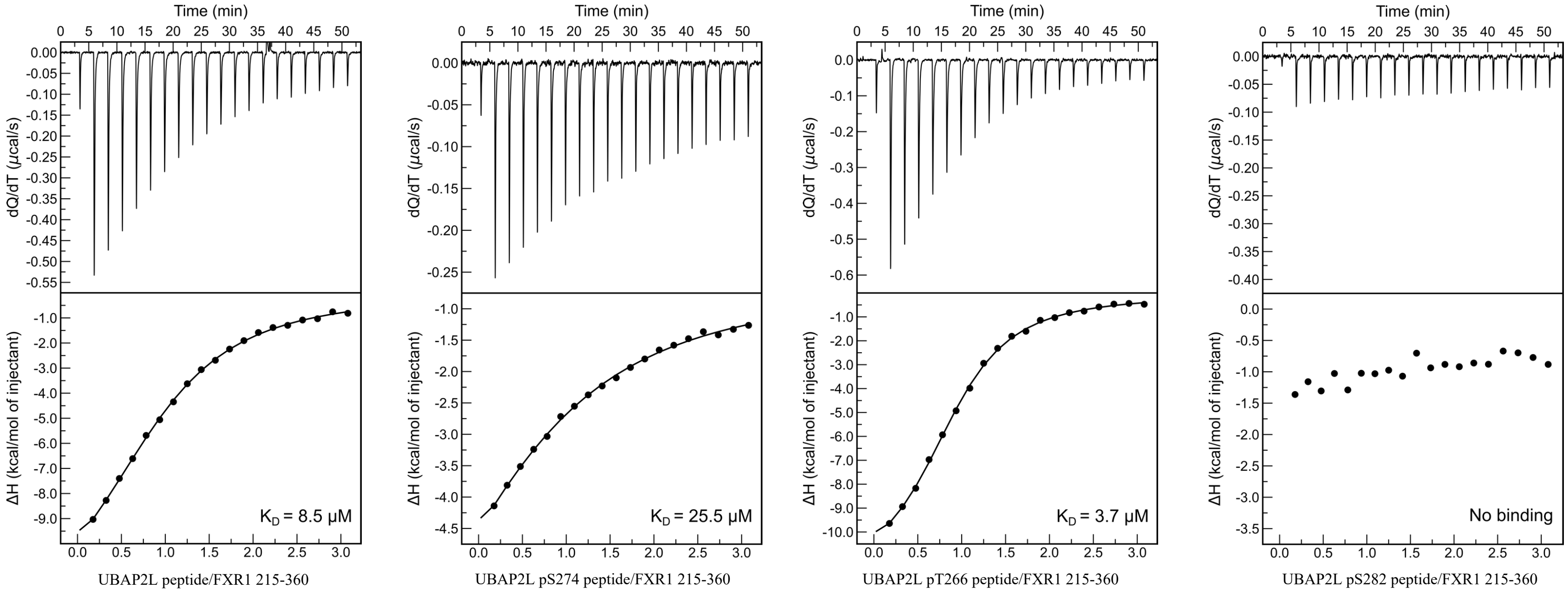


***Table 2 continued***

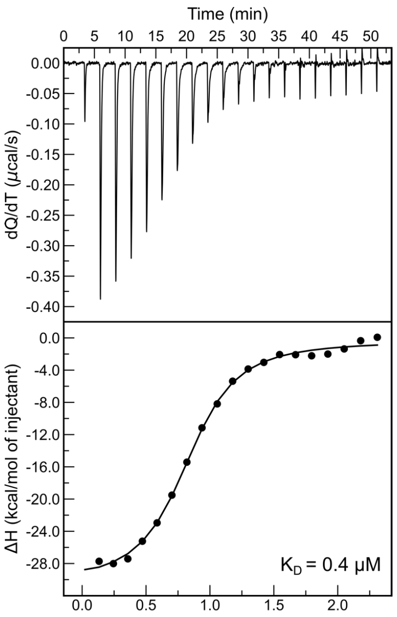


NF1 RNA/N protein SARS-CoV-2

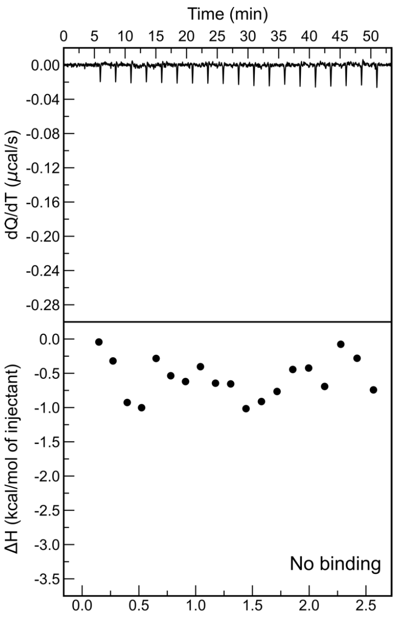


NF1 RNA/FXR1 215-360

*Table 2 continued*
